# Virulence priming of *Listeria monocytogenes* inside replication-permissive endolysosomes

**DOI:** 10.1101/2025.09.02.673654

**Authors:** Thomas J. P. Petit, Simon Guette-Marquet, Adrien Candat, Anouk Viry, Nicolas Desprat, Alice Lebreton

## Abstract

*Listeria monocytogenes* (*Lm*), the foodborne pathogen responsible for listeriosis, is well-known for its intracellular lifestyle. After entering epithelial cells, it can evade within minutes from its internalization vacuole into the cytosol, or transiently co-opt and replicate within endomembrane compartments termed epithelial Spacious *ListeriA*-containing Phagosomes (eSLAPs). Although eSLAPs display endolysosomal features, we bring evidence that they lack cathepsin D activity and are permeated. Indeed, sustained eSLAP membrane perforation by listeriolysin O is crucial for accommodating bacterial growth and vacuole expansion. Conversely, we show that *Lm* broad range phospholipase C PlcB is the main escape factor from eSLAPs. Unexpectedly, we find that *Lm* PrfA-dependent cytosolic virulence program is primed inside eSLAPs. As a consequence, the bacteria released from eSLAPs are pre-equipped with the actin assembly-inducing protein ActA and can recruit actin faster. We propose that a delayed access to the cytosol benefit eSLAP-transiting bacteria by favoring their dissemination or evasion from xenophagy.

## Introduction

*Listeria monocytogenes* (*Lm*) is a saprophytic bacterium and a zoonotic pathogen that can cause a foodborne infection known as listeriosis in wildlife, cattle and humans. While human listeriosis is rare, its typical symptoms (meningitis, encephalitis or sepsis) are severe for vulnerable individuals including immunocompromised patients, elderly persons or new-born babies, leading to a high fatality rate — 19.7% in the EU in 2023.^1^ Maternofetal transmission of *Lm* during pregnancy also remains a concerning cause of miscarriage, stillbirth, and preterm birth due to the high tropism of this bacterium for the placenta.^2,3^

*Lm* penetrates and thrives in mammalian hosts thanks to its ability to invade a wide range of host cell types, which allows it to cross epithelial or endothelial barriers, colonize tissues, and co-opt motile cells as vehicles throughout the organism.^4^ *Lm* predominantly resides and multiplies in the cytosol of its host cells, where recruitment of the actin cytoskeleton by the surface-exposed virulence factor ActA enables its motility, its capacity to spread from cell to cell, and confers a protection from xenophagy.^5–7^ Nonetheless, a subset of the bacterial population can also survive for periods ranging between minutes and days within several types of endomembrane compartments.^8,9^ Such alternative intracellular lifestyles, though long underappreciated, may significantly impact disease progression, bacterial fitness and shedding by providing niches for persistence, favoring immune or antibiotic avoidance, eliciting heterogeneity in the gene expression programs, or orienting dissemination strategies.^9,10^

The origin, maturation and properties of these compartments and the fate of the bacteria they contain differ between cell types. After its entry into goblet cells on the luminal side of intestinal villi, *Lm* can be transcytosed towards the basolateral side and released by exocytosis into the lamina propria in less than 30 min without leaving its internalization vacuole.^11^ In murine macrophages, *Lm* has been shown to reside and slowly replicate in non-acidic compartments termed spacious *Listeria*-containing phagosomes (SLAPs) that are decorated with the lysosomal marker LAMP1 and with the autophagy adapter LC3^12^. LC3-associated phagocytosis (LAP) is suggested to sort some of the bacteria towards this compartment during the early stage of macrophage invasion.^13^ In hepatocytes or trophoblasts infected for several days, cytosolic bacteria that gradually lose ActA on their surface become entrapped in acidic vacuoles decorated with lysosomal markers, independently of canonical autophagy.^14^ Persister bacteria were monitored for up to 25 days inside these *Listeria*-containing vacuoles (LisCVs); some of them entered a viable but non-culturable (VBNC) state and became tolerant to antibiotic treatments. *Lm* can also persist in a VBNC state in acidic, LAMP1-positive phagolysosomes in bovine polymorphonuclear neutrophiles before spreading to other cells and resuming a canonical life cycle.^15^

Real-time fluorescence monitoring of virulence factors during infection has proven well-suited to explore these alternative intracellular lives. It has previously led us to unveil another transient vacuolar niche where *Lm* could reside for several hours in epithelial cells, and which was termed eSLAPs by analogy with SLAPs.^16^ Both SLAPS and eSLAPs are decorated by LC3 and LAMP1 but non-acidic. Because the residence and growth of *Lm* inside either of these compartments depends on the secretion of the pore-forming toxin (PFT) listeriolysin O (LLO), the activity of this toxin is assumed to oppose their maturation into bactericidal phago– or endolysosomes, and potentially to allow the diffusion of cytosolic nutrient sources across the vacuole membrane. A major difference between SLAPs and eSLAPs is the bacterial growth rate. *Lm* multiplies inside eSLAPs as rapidly as in the cytosol (doubling time ∼1 h 20 min), whereas its doubling time is ∼8 h in SLAPs, suggesting that the epithelial intravacuolar niche is a less stringent environment. The rapid growth rate in eSLAPs also implies that their membrane needs to be continually expanded.

While eSLAPs have been shown to derive from primary internalization vacuoles,^16^ the bacterial and host cell mechanisms that drive the maturation of this niche into a compartment permissive for growth and the factors that govern the equilibrium between stability and rupture of the eSLAP membrane remained to be elucidated. *Lm* membrane-damaging virulence factors are anticipated to control the permeability of the vacuole membrane and the dynamics of escape towards the cytoplasm. In addition to LLO, the *Listeria* pathogenicity island-1 (LIPI-1) encodes two phospholipases: the phosphatidylinositide-specific phospholipase C PlcA, and the broad-range phospholipase C PlcB.^17^ Both have previously been shown to contribute to the escape of *Lm* from diverse types of endomembrane compartments, including internalization vacuoles, secondary vacuoles that are formed upon cell-to-cell spread, and pre-autophagosomes.^18,19^

The transcriptional activator PrfA is the central regulator of *Lm* virulence genes primordial for its intracellular life, including LIPI-I-encoded *hly*, *plcA*, *plcB* and *actA*.^20,21^ The efficiency of binding of PrfA to its target promoters is modulated by a conformational transition driven by the binding of its allosteric activator glutathione (GSH).^22,23^ In its unbound state, the affinity of PrfA for its targets is low; it can only induce the transcription of early virulence genes that harbor an optimal PrfA-binding sequence in their promoters, such as *hly* and *plcA*. Upon binding of GSH, the affinity of PrfA for its targets is enhanced, allowing the transcription of genes controlled by promoters harboring a more relaxed PrfA-binding site, such as the *actA-plcB* operon.^24–26^ Previous studies have led to a near-consensus that the allosteric transition of PrfA to its high affinity form occurs primarily upon *Lm* access to the host cell cytosol, where it sustains the expression of genes primordial for cytosolic colonization including *actA*, although a few reports have proposed that PrfA activation may also occur in phagosomes.^27–29^

To refine our understanding of the mechanisms that sustain the existence of eSLAPs and of their relevance during infection, we have set out to elucidate the ultrastructure of eSLAPs, the host and bacterial molecular mechanisms mediating their biogenesis, growth and disruption, and the virulence properties of the bacteria transiting through this route. Notably, we show that *Lm* co-opts the host endolysosomal system to establish eSLAPs in epithelial cell lines of intestinal, hepatic or mammary origin. We dissect the respective contributions of LLO and PlcB in the conversion of the internalization vacuole into a replicative niche, or in the destabilization of eSLAP membranes. Last, we identify that, in contrast to internalization vacuoles, *Lm* cytosolic virulence program is primed in eSLAPs, allowing a faster recruitment of actin upon release into the cytosol.

## Results

### Human epithelial cell lines of intestinal, mammary or hepatic origin are permissive for eSLAPs

We previously reported that *L. monocytogenes* can replicate within eSLAPs in LoVo and Caco-2 epithelial cells originating from colon adenocarcinoma.^16^ To gain further insight into the range of host cell types that are permissive for eSLAPs, we assessed eSLAP formation in eight other cell lines following infection with *Lm*. Cell lines were considered permissive if they harbored *Lm*-containing endomembrane compartments with typical eSLAPs features, *i.e*. which were positive for LAMP1 and LC3 in immunofluorescence staining (Fig S1A) and where bacterial replication was observed in live microscopy (Fig S1 and Movies S1 to S6). eSLAPs were detected in three additional colon cancer epithelial cell lines (LS174T, T84 and HCT-116), as well as in two breast adenocarcinoma cell lines (MCF7 and SK-BR-3) and in a hepatocyte-derived carcinoma cell line (Huh-7) (Fig S1 and Movies S1 to S6), indicating that this vacuolar niche is not restricted to colonocytes. By contrast, no growth of intravacuolar bacteria was observed after infection of the lung adenocarcinoma cell line Calu-3 or of the placental choriocarcinoma cell line JEG-3 by *Lm*.

### eSLAPs interact with the host cell endolysosomal system

The endomembrane compartments characterized as eSLAPs are decorated with markers of the endolysosomal maturation pathway, including Rab7 and LAMP1.^16^ While LAMP1 is commonly used as a marker of late endosomes and lysosomes, it was also shown to be recruited at the *Salmonella*-containing vacuole by a process independent of lysosome fusion.^30,31^ To gain a deeper understanding of eSLAP maturation and interactions with the host cell trafficking, we assessed the colocalization of several other endolysosomal markers with eSLAPs. Immunofluorescence staining of LoVo cells infected with *Lm* for 3 h revealed the presence of ATP6V1A, a subunit of the vacuolar-type ATPase (v-ATPase) complex, of LIMP-2, another lysosomal membrane protein, and of the lysosomal protease cathepsin D around DAPI-stained bacteria gathered inside eSLAPs (Fig 1A). By contrast, in live-microscopy experiments using SiR-Lysosome —a fluorogeneic probe that reports on the enzymatic activity of cathepsin D— in infected LoVo cells, no signal was detected inside eSLAPs (Fig 1B). Punctiform SiR-Lysosome staining was decorating the immediate vicinity of eSLAPs, which may indicate the presence of lysosomes interacting with the pathogen-containing vacuole prior to fusion. Inside eSLAPs, Cathepsin D thus appears to be present but inactive. Given an optimum pH of 5 for this enzyme,^32^ this lack of activity is consistent with our previous observation that eSLAPs are not acidic.^16^ Altogether, these observations suggest that although eSLAPs display several features of late endosomes/lysosomes, they fail to fully acquire degradative characteristics.

**Fig 1.**
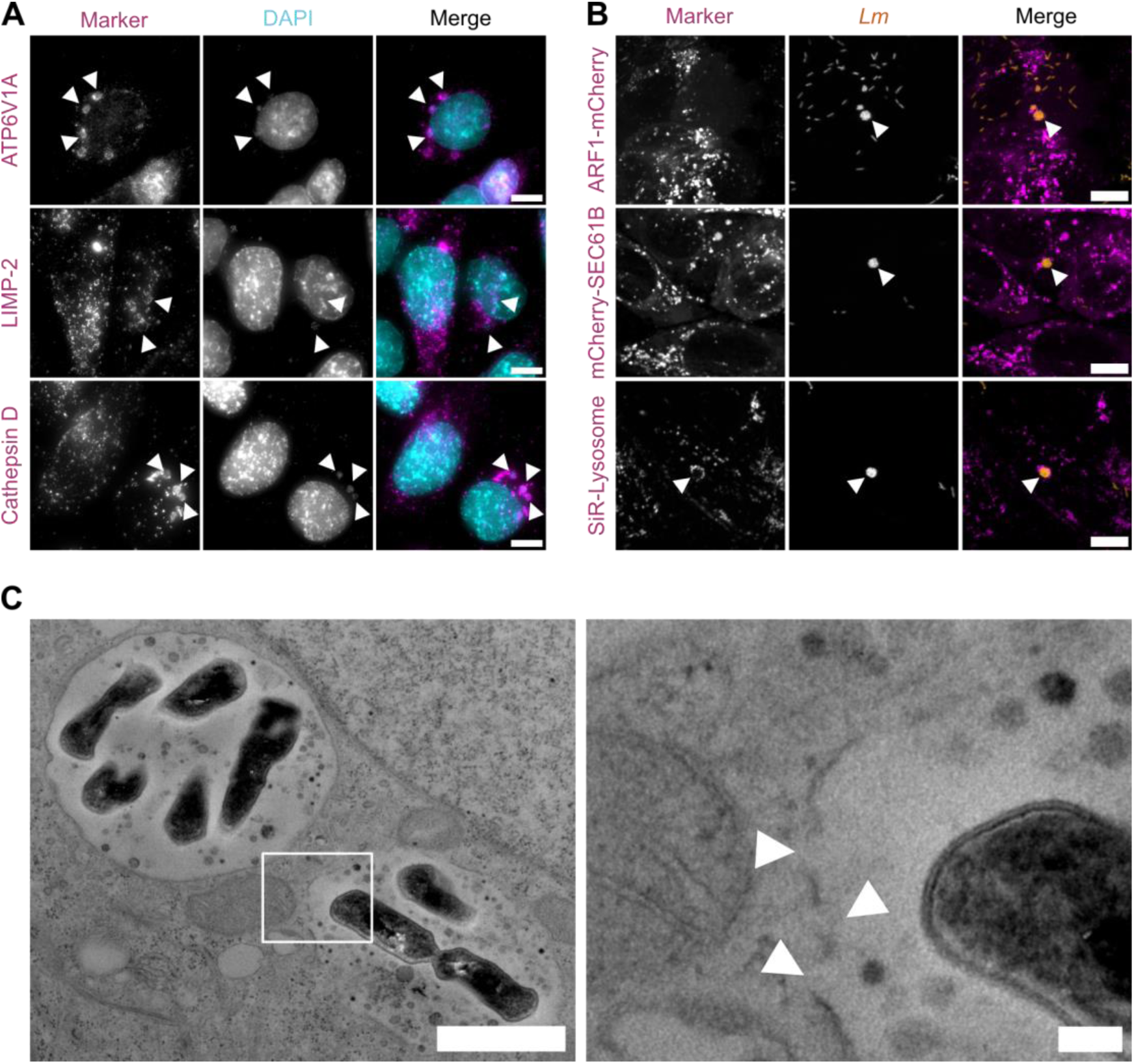
eSLAPs interact with endolysosomal compartments and exhibit membrane permeabilization. (A) Images of immunofluorescence targeting ATP6V1A, LIMP-2 and Cathepsin D (in magenta) in LoVo cells infected by *Lm* (stained with DAPI, in cyan) at a MOI of 100 for 3 h. The white arrowheads point to representative eSLAPs labelled with the associated marker. Scale bars: 10 μm. (B) Spinning disk microscopy images of live LoVo cells expressing ARF1-mCherry, mCherry-SEC61B or stained with SiR-Lysosome (in magenta) infected with *Lm* expressing mCherry (in orange) at 3 hours post-infection (hpi) at a MOI of 20. The white arrowheads point to representative eSLAPs. Scale bars: 10 μm. (C) Representative eSLAPs in LoVo cells infected by *Lm* for 3 h (MOI 100) observed by TEM at two magnifications. Scale bars: 1 μm (left), 100 nm (right). The white arrowheads point to eSLAP membrane discontinuities.

To evaluate the contribution of other organelles to eSLAP biogenesis, we monitored the localization of the endoplasmic reticulum (ER) marker mCherry-SEC61B and Golgi apparatus marker ARF1-mCherry expressed by live microscopy in LoVo cells stably expressing these constructs and infected with *Lm* expressing GFP. Neither marker was detected at the eSLAP membrane, suggesting an absence of trafficking between the ER or Golgi apparatus and eSLAPs (Fig 1B).

### eSLAPs are delimited by a single membrane with local discontinuities

To gain insight into the ultrastructure of eSLAPs, we performed transmission electron microscopy (TEM) on LoVo cells infected for 3 h with *Lm*. eSLAPs appeared as large, single-membrane vacuolar compartments containing dividing bacteria (Fig 1C). Within the lumen of these compartments, numerous electron-dense vesicle-like structures of 20-100 nm in diameter were observed, which could resemble intraluminal vesicles typically found in multivesicular bodies (MVBs). By places, eSLAPs displayed membrane discontinuities, indicating that these compartments are eventually permeated. The requirement of *Lm*-secreted listeriolysin O for bacterial growth in eSLAP,^16^ their neutral pH and poor bactericidal activity contrasting with their otherwise endolysosomal characteristics, are consistent with their membrane being at least transiently damaged.

### Markers of endomembrane damage are continuously recruited at the eSLAP membrane

Given the observation of membrane discontinuities in eSLAPs by TEM, and considering that eSLAPs can grow and persist several hours without rupture, we hypothesized that eSLAPs might either remain intact until shortly before rupture, or exist in a constitutively permeated state. To investigate the dynamics of eSLAP membrane permeation, we monitored the recruitment of galectin-3 (Gal3) and galectin-8 (Gal8), two host lectins that accumulate on damaged endomembranes by recognizing exposed glycans.^33^ LoVo cells stably expressing mAG-Gal3 or EGFP-Gal8 were infected with *Lm* expressing mCherry, then eSLAPs were imaged by spinning disk live-cell microcopy (Fig 2A and 2C, Movies S7 and S8). Fluorescence measurement at the position of eSLAPs revealed that both galectins were recruited at eSLAP membranes from the earliest time-points of observation and remained associated with eSLAPs throughout their lifespan (Fig 2B and 2D). Upon eSLAP membrane breakdown and bacterial escape, the intensity of mAG-Gal3 or EGFP-Gal8 fluorescence signals increased abruptly, marking a more massive recruitment of galectins at the disassembly stage of the vacuole than during the phase of bacterial residence. These findings suggest that eSLAPs are chronically permeated compartments that continuously expose luminal glycans to the cytosol.

**Fig 2.**
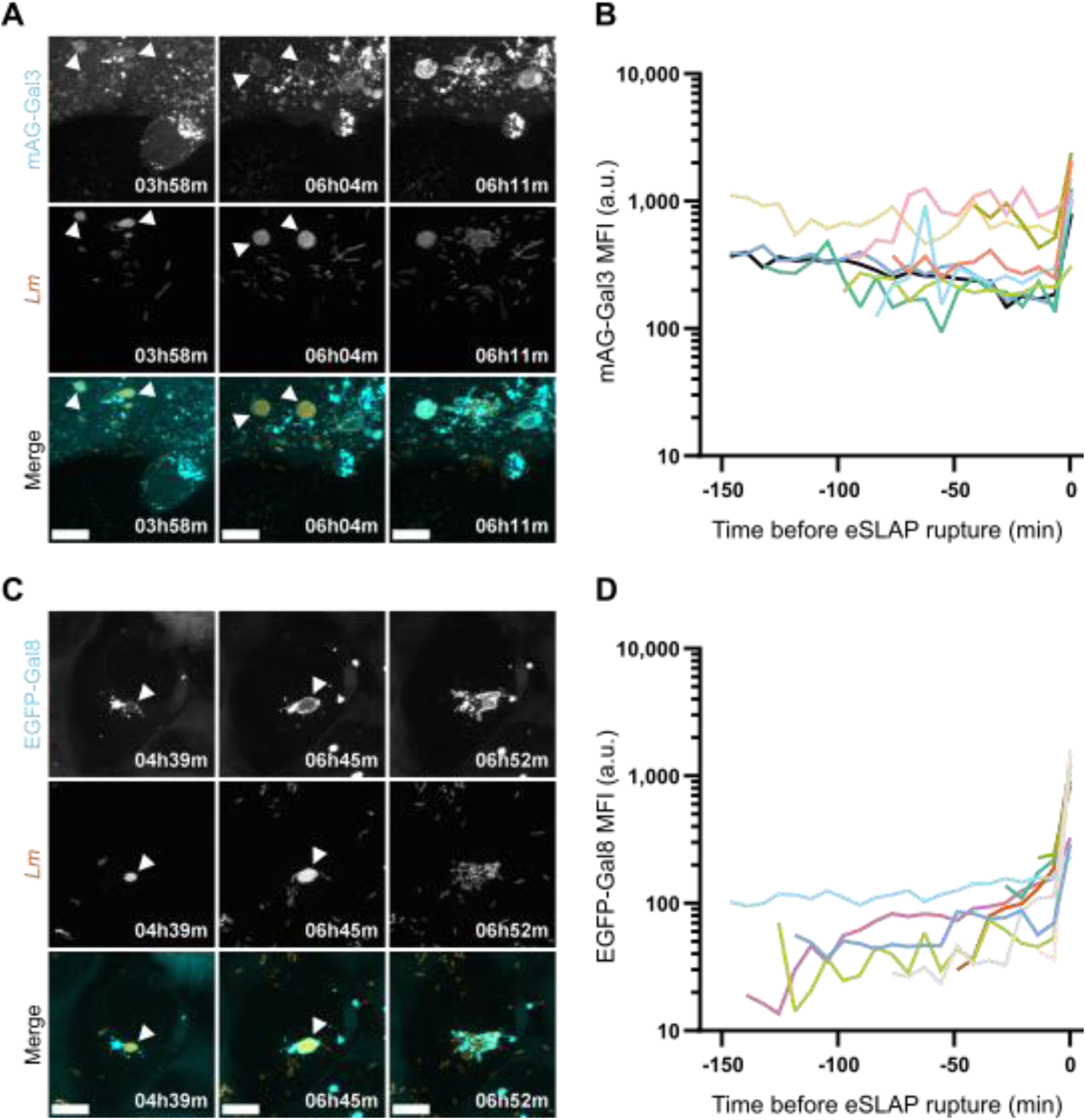
The membrane damage markers Gal3 and Gal8 are continuously recruited at the eSLAP membrane. (A, C) Representative live spinning disk microscopy images of LoVo cells expressing mAG-Gal3 (A) or EGFP-Gal8 (C) (cyan) infected with *Lm* expressing mCherry (orange, MOI 20) at several time points post-infection. The white arrowheads point to representative eSLAPs and the associated marker. Scale bars: 10 μm. (B, D) Quantification of the mean fluorescence intensity (MFI) of mAG-Gal3 (B) and EGFP-Gal8 (D) colocalizing with eSLAPs over time, relatively to the time of eSLAP rupture taken as a common reference time (t = 0 min). Each colored line represents the MFI quantification for one eSLAP (n = 9 for mAG-Gal3, n = 10 for EGFP-Gal8). Data are displayed on a logarithmic scale on the Y axis.

### The pore-forming activity of LLO is required for *Lm* replication in eSLAPs

*Lm* secretes several virulence factors with membrane-damaging properties such as phospholipases and the pore-forming toxin LLO. We have previously observed that a Δ*hly* mutant that does not produce LLO cannot replicate in eSLAPs.^16^ However, a strain that secretes a mutant version of LLO (W_492_A) with low residual activity can still replicate in these compartments. Whereas we had previously estimated the residual activity of LLO_W492A_ between 1 and 3 % by assessment on murine erythrocytes,^16^ reproducing this assay on horse erythrocytes revealed that this version of LLO retained 16 ± 6 % of activity compared to wild type (WT) LLO (Table S4). Considering this information, and given that LLO can modulate other cellular functions independently of its pore-forming activity, notably as a pathogen-associated molecular pattern (PAMP),^34–36^ it remained unclear whether the pore-forming activity of LLO itself was required for *Lm* replication in eSLAPs. To clarify its role, we generated a strain secreting a detoxified form of LLO, dtLLO (Fig S2A),^34^ that exhibited a 97 ± 2 % reduction in hemolytic titer on horse erythrocytes compared to the WT strain (Fig 3A and Fig S2B). LoVo cells were infected for 3 h with *Lm* strains expressing mCherry and either producing WT LLO, lacking LLO (Δ*hly*) or producing dtLLO (dt*hly*), after which eSLAP frequency was quantified for each strain. The ability of both the Δ*hly* and dt*hly* strains to generate eSLAP-like compartments was severely compromised compared to the WT strain (Fig 3B and 3C), to an extent that cannot be attributed to the modest impact (∼25% reduction) of these two mutations on bacterial invasion alone (Fig S2C). These results confirm that the pore-forming activity of LLO constitutes a critical determinant for *Lm* replication within eSLAPs.

**Fig 3.**
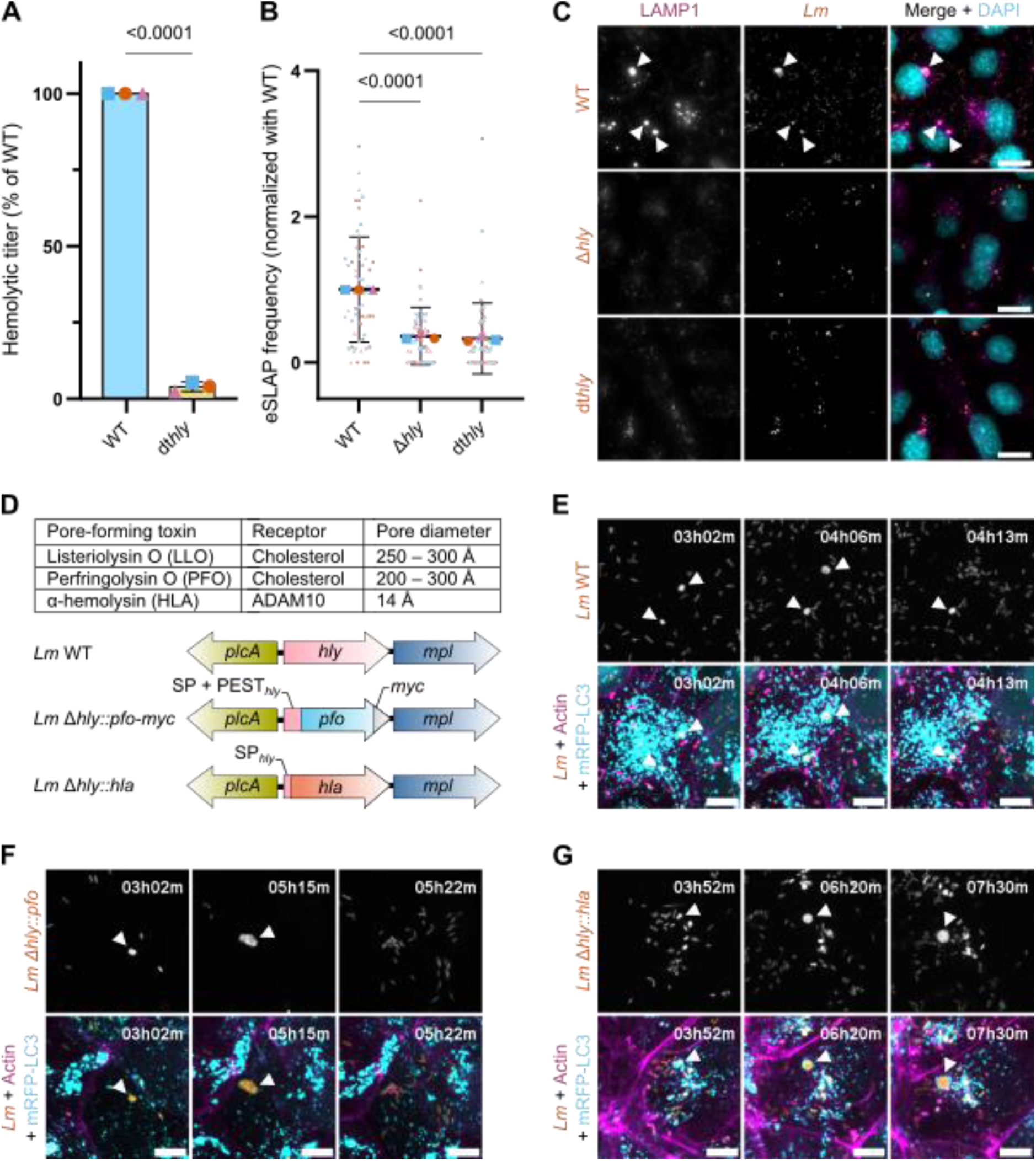
Bacterial growth within eSLAPs depends on a pore-forming activity. (A) Hemolytic properties of the *Lm* strains expressing WT LLO or detoxified LLO variant C_484_A-W_491_A-W_492_A (dt*hly*). The average hemolytic titers and SD of three independent experiments are represented. The *p*-value indicates the result of a two-tailed Student’s *t*-test on the distributions, assuming equal variance. (B) Quantification of eSLAP frequency in LoVo cells infected for 3 h by *Lm* WT, dt*hly* and Δ*hly* (MOI 100), all expressing mCherry. Statistical comparison was done with a Kruskal-Wallis test with Dunn’s multiple comparisons, assuming equal variance. Data from three independent experiments (each one of them indicated with a different color) are represented in “SuperPlots”^64^ where each value of eSLAP frequency per image is shown (small symbols) together with the means of each experiment (larger symbols). The mean ± SD from all data are indicated in black. (C) Representative immunofluorescence images of the infections quantified in (B). The white arrowheads point to representative eSLAPs (*Lm*, orange) decorated by the marker LAMP1 (magenta). DAPI was used to visualize nuclei (cyan). Scale bars: 20 μm. (D) Table summarizing the main properties of the different pore-forming toxins (PFTs) —LLO, PFO and HLA— used in this study. The genomic locus in which *hly* was replaced with SP+PEST*_hly_*-*pfo-myc* (*Lm* Δ*hly::pfo*) or SP*_hly_*-*hla* (*Lm* Δ*hly::hla*) is represented underneath the table. (E-G) Representative live spinning disk microscopy images of LoVo cells expressing mRFP-LC3 (cyan) infected with *Lm* WT at MOI 20 (E), *Lm* Δ*hly::pfo* at MOI 30 (F), and *Lm* Δ*hly::hla* at MOI 50 (G) all expressing GFP (orange) at several time points post-infection. The white arrowheads point to representative eSLAPs labelled with mRFP-LC3. F-actin is shown in magenta. Scale bars: 10 μm.

### eSLAP permeation allows *Lm* replication in eSLAPs independently of the nature of the pore-forming toxin employed

To further probe the influence of membrane perforation on the existence of the eSLAP replicative niche, we generated *Lm* strains in which the *hly* gene within the LIPI-I was replaced with genes encoding heterologous PFTs (Fig 3D). Purposedly, we replaced *hly* with the gene encoding perfringolysin O, another member of the cholesterol-dependent cytolysins (CDC) family closely related to LLO. The PEST sequence of *hly* was kept in the construct to decrease PFO cytotoxicity,^37^ giving rise to the strain *Lm* Δ*hly*::PEST*_hly_*-*pfo*-*myc*. In *Lm* Δ*hly*::SP*_hly_*-*hla*, *hly* was replaced with the gene encoding the α-hemolysin (HLA) of *Staphylococcus aureus*, a more distantly related PFT that generates smaller pores than CDCs (Fig 3D). The secretion of HLA by *Lm* was allowed through its N-terminal fusion with the *hly* signal peptide. The engineered strains expressing HLA and PFO secreted the toxins as expected (Fig S2D). The strain secreting PFO exhibited a hemolytic activity comparable to that of the WT strain, indicating functional activity of the toxin (Fig S2E). All strains additionally expressed constitutively GFP for live-cell microscopy tracking experiments.

In order to address whether the pore-forming activities of these PFTs could substitute for LLO in supporting *Lm* replication within eSLAPs, we infected LoVo cells expressing mRFP-LC3 with the strains producing each PFT and monitored the intravacuolar growth of bacteria by live-cell microscopy. Remarkably, both the PFO and HLA-producing strains were capable of replication in LC3-labelled compartments similar to the ones observed with the WT strain (Fig 3E to 3G, Movies S9 to S11). PFTs can thus sustain *Lm* replication in vacuoles regardless of the protein scaffold that allows the formation of the pore and of its diameter. Hence, these results reinforce the key role endorsed by the pore-forming activity of secreted cytolysins for the existence of eSLAPs.

### The broad range phospholipase C PlcB destabilizes eSLAPs

*Lm* secretes two phospholipases C, PlcA and PlcB, that have been implicated in several steps of the infection cycle including escape from the internalization vacuole (PlcA) and from the secondary vacuole after cell-to-cell spread (both PlcA and PlcB).^18^ Given the membrane-destabilizing properties of these virulence factors, we sought to determine whether they also contributed to eSLAP dynamics. To this end, *Lm* mutants lacking the sequence encoding PlcA or PlcB (*Lm* Δ*plcA* and *Lm* Δ*plcB*, respectively), both expressing mCherry, were generated. At 3 hpi infection in LoVo cells, a 3-fold increase in eSLAP frequency was quantified for *Lm* Δ*plcB* compared to the WT strain (Fig 4A and 4B), whereas deletion of *plcA* had no significant effect on eSLAP frequency (Fig S3A). Infection of LoVo cells with a complemented Δ*plcB* strain that harbors an integrative plasmid with *plcB* under the control of its native promoter P*_actA_* restored eSLAP frequency to WT levels (Fig 4A and 4B, Fig S3B). Importantly, the increase in eSLAP frequency measured in absence of PlcB was not attributable to effects on bacterial entry or intracellular replication, as the number of bacteria per cell at 1 hpi and 3 hpi was similar across all strains (Fig S3C and S3D). Given the known enzymatic activity of PlcB as a phospholipase, the most likely explanation for its impact on eSLAP frequency is that it promotes the destabilization of eSLAP membranes and favors bacterial escape into the cytosol.

**Fig 4.**
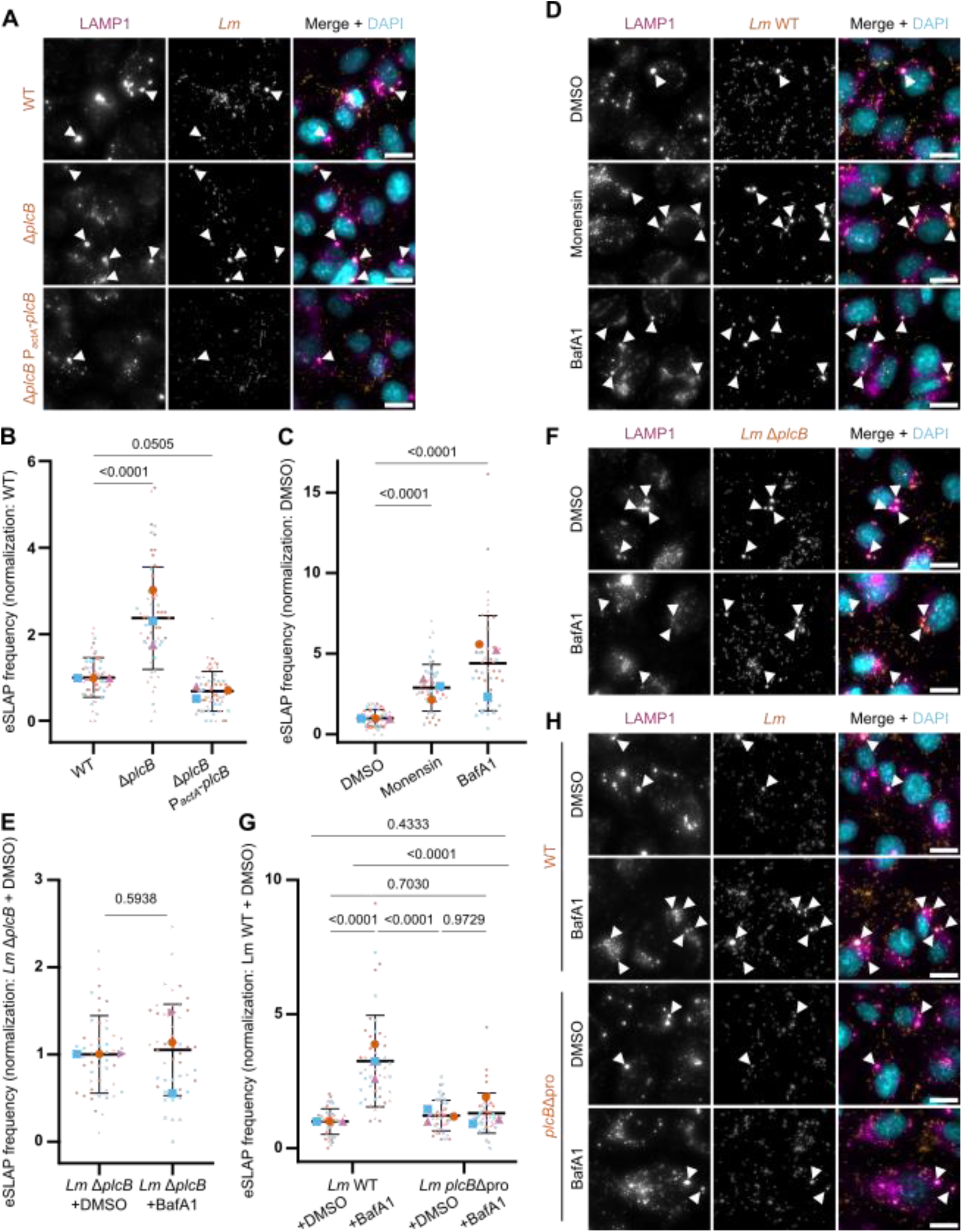
Dependance of eSLAPs on the acid-dependent maturation of PlcB. (A) Representative immunofluorescence images of LoVo cells infected for 3 h by *Lm* WT, Δ*plcB* and Δ*plcB*-P*_actA_*-*plcB* (MOI 100), all expressing mCherry. the infections quantified in (B). The white arrowheads point to representative eSLAPs (*Lm*, orange) decorated by the marker LAMP1 (magenta). DAPI was used to visualize nuclei (cyan). Scale bars: 20 μm. (B) Quantification of eSLAP frequency in LoVo cells infected as indicated in (A). (C) Quantification of eSLAP frequency in LoVo cells infected for 3 h by *Lm* WT expressing mCherry (MOI 100), and treated with DMSO, monensin (100 μM) or BafA1 (100 nM) during the first hour of infection. (D) Representative immunofluorescence images of the infections quantified in (B). The white arrowheads point to representative eSLAPs (*Lm* WT, orange) decorated by the marker LAMP1 (magenta). DAPI was used to visualize nuclei (cyan). Scale bars: 20 μm. (E) Quantification of eSLAP frequency in LoVo cells infected for 3 h by *Lm* Δ*plcB* expressing mCherry (MOI 100), and treated with DMSO or BafA1 (100 nM) during the first hour of infection. (F) Representative immunofluorescence images of the infections quantified in (D). The white arrowheads point to representative eSLAPs (*Lm* Δ*plcB*, orange) decorated by the marker LAMP1 (magenta). DAPI was used to visualize nuclei (cyan). Scale bar: 20 μm. (G) Quantification of eSLAP frequency in LoVo cells infected for 3 h by *Lm* WT or *Lm plcB*Δpro expressing mCherry (MOI 100), and treated with DMSO or BafA1 (100 nM) during the first hour of infection. The *p*-values indicate the results of a two-way ANOVA with, assuming equal variance. (H) Representative immunofluorescence images of the infections quantified in (F). The white arrowheads point to representative eSLAPs (*Lm* WT or *Lm plcB*Δpro, orange) decorated by the marker LAMP1 (magenta). DAPI was used to visualize nuclei (cyan). Scale bar: 20 μm. For (B, C, E, and G), the mean ± SD from three independent experiments is displayed. For (B, D, F), each color corresponds to one biological replicate. The *p*-values indicate the results of one-way ANOVAs with Dunnett’s correction for multiple comparisons in (B, C and E), or Šídák’s correction for multiple comparison in G, assuming equal variance.

### Impeding vacuole acidification favours eSLAPs

As indicated above, eSLAPs are decorated with ATP6V1A, a subunit of the v-ATPase complex (Fig 1A). Because this complex is driving the acidification of endomembrane compartments through its activity as a proton pump, we questioned whether it was contributing to the dynamics of eSLAPs, although we had not observed their acidification.^16^ To gain insight into the effects of host cell endolysosomal acidification mechanisms on the occurrence of eSLAPs, we assessed the impact of two pharmacological treatments on eSLAP frequency, by exposing cells either to 100 μM of monensin or to 100 nm of bafilomycin A1 during the first hour of infection. Monensin acts as an ionophore that can dissipate proton gradients across endomembranes, while bafilomycin A1 directly interferes with the activity of the v-ATPase. There was no impact of either treatment on the invasion of LoVo cells by bacteria (Fig S3E). By contrast, both treatments significantly increased the frequency of eSLAPs quantified at 3 hpi, with respectively a 2.9-fold increase for monensin, and a 4.4-fold increase for bafilomycin A1 (Fig 4C and 4D). These results suggest that v-ATPase-dependent mechanisms of acidification of endomembrane compartments contribute either to restricting the growth of *Lm* inside eSLAPs or to the stability of these compartments.

### Control of eSLAP acidification is gating PlcB-dependent escape

Because eSLAPs are poorly acidic and negative for cathepsin activity even in absence of monensin or bafilomycin, we reasoned that dampening of intravacuolar toxicity was unlikely to explain the strong promoting effect of these drugs on eSLAP frequency. We thus hypothesized that by opposing moderate or transient events of vacuole acidification, these drugs have the potential to interfere with the activity of membrane-damaging effectors and thereby increase the frequency of eSLAPs measured at a given time-point by reducing the probability of membrane rupture. PlcB stood as a promising candidate as an indirect target of these drugs: as we have seen above, it is involved in the destabilisation of eSLAPs, and bafilomycin treatment phenocopies the deletion of *plcB* with regard to the frequency of eSLAPs (Fig 4B and 4C). Moreover, the activity of PlcB is known to be controlled by pH. Indeed, PlcB is initially synthesised and secreted by Sec as an inactive proenzyme (Pro-PlcB) that is retained at neutral pH between the bacterial membrane and cell wall due to a N-terminal propeptide.^38^ PlcB maturation depends on its companion metalloprotease Mpl.^39^ In acidic conditions, Mpl acquires its proteolytic activity by autocatalytic cleavage, and then cleave the retained pool of Pro-PlcB into mature PlcB.^40^ Mature PlcB is then released from the bacterial surface, which allows it to reach its substrates in the vacuole membrane.^38,40^

To infer whether the effect of bafilomycin on eSLAP occurrence was mediated through an impact on PlcB maturation, we assessed eSLAP frequency in conditions where we combined bafilomycin treatment with the use of either a Δ*plcB* strain, or a *plcB*Δpro strain. In the latter, the sequence of the PlcB propeptide was deleted, thus bypassing the need for pH-dependent maturation. For both strains, eSLAP frequency became insensitive to bafilomycin: it was maximized with the Δ*plcB* strain (Fig 4E and 4F), and minimized with the *plcB*Δpro strain (Fig 4G and 4H). These two complementary results indicate that the positive effect on eSLAP frequency observed when interfering with the acidification of endomembrane compartments is primarily mediated by delaying on PlcB maturation and release. Altogether, our findings position mature PlcB as the main escape factor from eSLAPs.

### The virulence master regulator PrfA is activated in *Lm* replicating in eSLAPs

Whereas PrfA is well-known to be allosterically-induced once *Lm* gains access to the cytosol, its status in eSLAPs was unknown. Yet, the reliance on PlcB activity for escape from eSLAPs suggested that the expression of *plcB*, which depends on the active form of PrfA, would be initiated in eSLAPs. To investigate the activation state of PrfA in bacteria replicating in eSLAPs, we engineered a fluorescent reporter strain constitutively expressing mCherry from an inert locus and carrying a *gfpmut2* gene encoding GFP under control of P*_actA_*, which is strictly dependent on PrfA activation (Fig 5A). To validate this reporter, LoVo cells were infected with this strain and monitored at two different time points: 25 min post-infection, when bacteria are still located within internalization vacuoles and when PrfA is expected to be weakly active, and 3 hpi, when most bacteria have reached the cytosol and when PrfA is expected to be highly active. As anticipated, no GFP-positive *Lm* were observed at 25 min post-infection, while GFP fluorescence was detected in cytosolic bacteria at 3 hpi, confirming the functionality and specificity of this reporter system (Fig 5B). We next assessed PrfA activation in the population of bacteria residing in eSLAPs at 3 hpi. At this stage, 95 % of eSLAPs contained GFP-positive bacteria, indicating that PrfA is in its active state within these vacuolar compartments (Fig 5B and 5C). Immunofluorescence targeting ActA further confirmed these results, as ActA staining colocalized with bacteria in 92 % of eSLAPs (Fig 5B and 5C).

**Fig 5.**
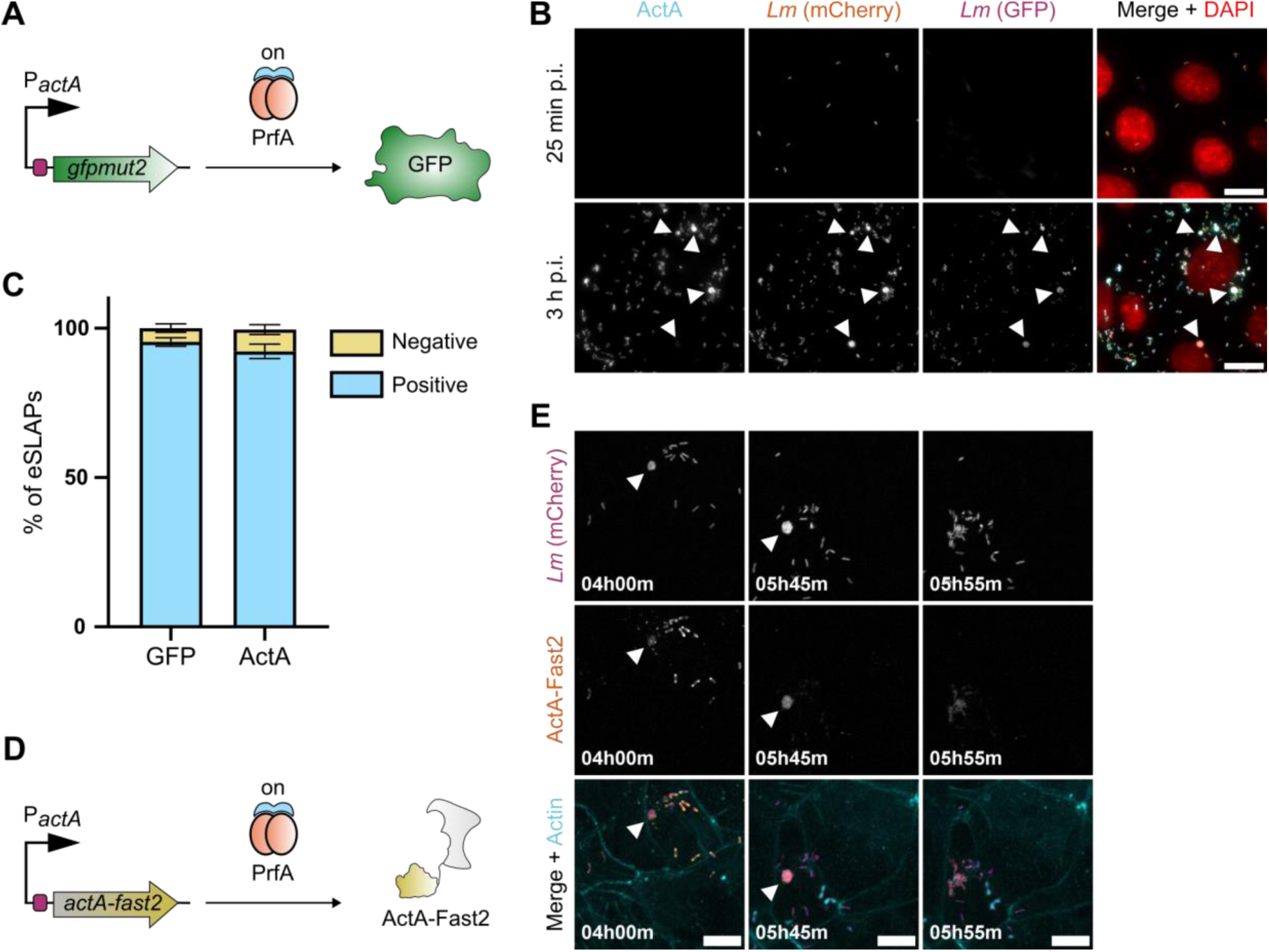
The virulence program controlled by PrfA is activated by bacteria residing in eSLAPs. (A) Diagram of the transcriptional reporter of PrfA activation, where expression of *gfpmut2* (encoding GFP) is under control of the strictly PrfA-dependent promoter P*_actA_* in the pAD vector. This construct was integrated in a strain of *Lm* that constitutively expresses mCherry under control of the P_HYPER_ promoter (*Lm* mCherry). (B) Representative images of LoVo cells infected by *Lm* mCherry pAD-P*_actA_*-*gfpmut2* (MOI 100) for 25 min and 3 h, and stained by immunofluorescence against ActA (cyan). At 3 h, the white arrowheads point to representative eSLAPs containing *Lm* (mCherry, orange) with activated PrfA (GFP, magenta) and decorated by ActA (cyan). DAPI was used to visualize nuclei (red). Scale bar: 20 μm. (C) Quantification of the proportion of eSLAPs that were positive (blue) or negative (yellow) for GFP fluorescence or ActA immunostaining in LoVo cells infected for 3 h with *Lm* mCherry pAD-P*_actA_*-*gfpmut2*. Data correspond to the mean ± SD from three independent experiments. (D) Diagram of the translational fusion construct where the expression of the *actA-fast2* fusion is under control of the strictly PrfA-dependent promoter P*_actA_* in the pAD vector. This construct was integrated in the constitutively-expressing mCherry strain *Lm* mCherry, (E) Representative live spinning disk microscopy images of LoVo cell infected with *Lm* mCherry pAD-P*_actA_*-*actA*-*fast2* (MOI 20) at several time points post-infection. The white arrowheads point to representative eSLAPs with *Lm* (mCherry, magenta) synthesizing ActA-Fast2 (orange). F-actin is shown in cyan. Scale bars: 10 μm.

To complement these observations by the direct visualization of ActA synthesis in live infected cells, we generated a strain carrying a translational fusion of ActA to the FAST2 tag, under the control of P*_actA_* (Fig 5D). As previously shown for FAST,^16^ FAST2 can be secreted by bacteria and becomes fluorescent upon addition of a fluorogenic chromophore, providing a live readout of ActA synthesis. Infection of LoVo cells with this strain and live-cell imaging revealed FAST2 fluorescence in eSLAPs where bacteria replicated, confirming active synthesis of ActA in these compartments (Fig 5E and Movie S12). Together, these results demonstrate that PrfA is fully activated in *Lm* replicating within eSLAPs and that virulence factors important for cytosolic colonization, such as ActA, are already produced by bacteria while still enclosed within this vacuolar niche.

### Prior residence in eSLAPs accelerates actin recruitment by *Lm* upon escape from vacuoles

Given that PrfA is highly active in eSLAPs and that virulence factors such as ActA are already synthetized by *Lm* replicating therein, we hypothesized that these vacuolar compartments could act as virulence priming niches, where the bacteria would acquire some of the molecular equipment later used for efficient cytosolic colonization. One benefit of such priming would be a faster recruitment of actin upon cytosolic access, compared to bacteria that escape early from internalization vacuoles.

To assess if the timing of actin recruitment differs between the two scenarios, we generated LoVo cells stably expressing EYFP fused to the cell-wall binding domain of listeriophage A500 (EYFP-CBD_500_), which labels *Lm* only when they are released into the cytosol of host cells.^41,42^ These cells were infected with *Lm* expressing mCherry and treated with SPY650-FastAct, a fluorescent actin probe, allowing real-time visualization of both bacterial access to the cytosol and actin recruitment by live-cell microscopy. The temporal difference between EYFP-CBD_500_ acquisition and actin recruitment was measured for individual bacteria as a proxy for the delay between vacuole rupture and actin polymerization (hereafter actin recruitment time) (Fig 6A). After escape from the internalization vacuole, bacteria were found to recruit actin with a median time of 25 min (Fig 6B and 6C, Movie S13), consistent with previous assessments of the respective timings of bacterial contact with the cytosol and of actin polymerization initiation.^42^ By contrast, bacteria escaping from eSLAPs recruited actin significantly faster than those escaping from internalization vacuoles, with a median recruitment time of 10 min (Fig 6B and 6C, Movie S14). While both populations contained a subset of bacteria for which no actin recruitment was detected over the 45 min of observation, the distribution of recruitment times was clearly shifted toward shorter delays for bacteria exiting eSLAPs. These findings support the hypothesis that eSLAPs behave as virulence priming niches, enabling *Lm* to synthetize key virulence factors while confined in a vacuolar environment and protected from cytosolic surveillance mechanism, and then to recruit actin more rapidly upon cytosolic access. This “bootcamp” strategy could confer advantages in evading host defenses such as xenophagy and promoting efficient cell-to-cell spread.

**Fig 6.**
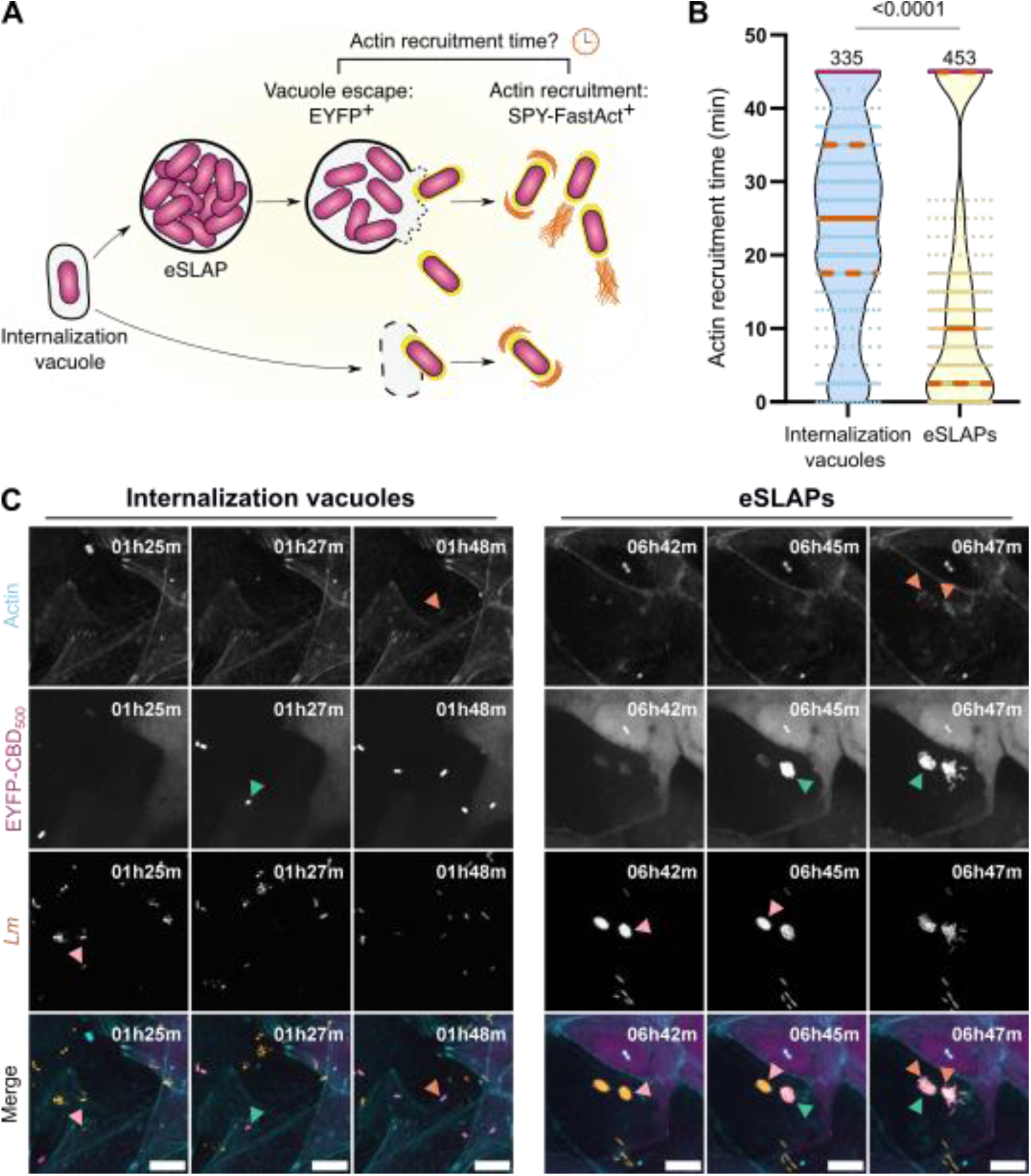
Residence in eSLAPs allows a rapid actin recruitment upon *Lm* escape into the cytosol. (A) Schematic representation of the method used to determine the timing of actin recruitment by bacteria after vacuole escape. Bacteria that reach the cytosol acquire the fluorescent-tagged cell wall binding domain EYFP-CBD_500_, and actin recruitment around bacteria can be detected with SPY650-FastAct staining. The actin recruitment time corresponds to the interval between the two time points when (*i*) EYFP is first detected and (*ii*) actin is first detected around the same bacterium. (B) Violin plots comparing actin recruitment time between bacteria escaping from internalization vacuoles or from eSLAPs, in LoVo cells infected with *Lm* expressing mCherry (MOI 20). Quartiles are represented by orange dotted lines and the median by a solid orange line. In both populations, bacteria that had not recruited actin within 45 min are plotted as purple dots in the upper limit of the distribution. In total, 335 bacteria escaping from internalization vacuoles and 453 bacteria escaping from eSLAPs (from 35 imaged eSLAPs) were tracked. The *p*-value represents the result of a Mann-Whitney *U* test to compare the two distributions. (C) Representative live spinning disk microscopy images of LoVo cells expressing EYFP-CBD_500_ (magenta) infected with *Lm* WT expressing mCherry (orange) and escaping from the internalization vacuole (left) or from eSLAPs (right). Pink arrowheads point at bacteria contained within a vacuole, green arrowheads point at bacteria that reached the cytosol attested by EYFP staining, and orange arrowheads point at bacteria that recruited actin (cyan). Scale bars: 10 μm.

## Discussion

In recent years *Lm* has emerged as a one of the invasive bacteria that can alternate between the two scenarios of intracellular life — being cytosolic or vacuolar — in a variety of cell types.^8,9^ However, our understanding of how this multiplicity of strategies may provide local benefits to a part of the bacterial population and subsequently alter the residence and proliferation of bacteria in tissues or their propagation remains mostly theoretical. Here, taking the example of the eSLAP niche that can transiently establish in intestinal, hepatic or mammary epithelial cells, we provide evidence that delayed residence inside endomembrane compartments results in a priming of *Lm* cytosolic virulence program that hastens actin recruitment and cytosolic motility once the bacteria eventually reach the cytosol. Our work also brings a refined picture of the interplay between bacterial and host factors that control the dynamics of eSLAP maturation, growth and rupture. Besides the dominant endolysosomal characteristics of these compartments, two bacterial effectors targeting endomembranes, LLO and PlcB, play contrasting roles in their permeation and stability. An overview of the main features of eSLAP dynamics highlighted in this study is illustrated in Fig 7.

**Fig 7.**
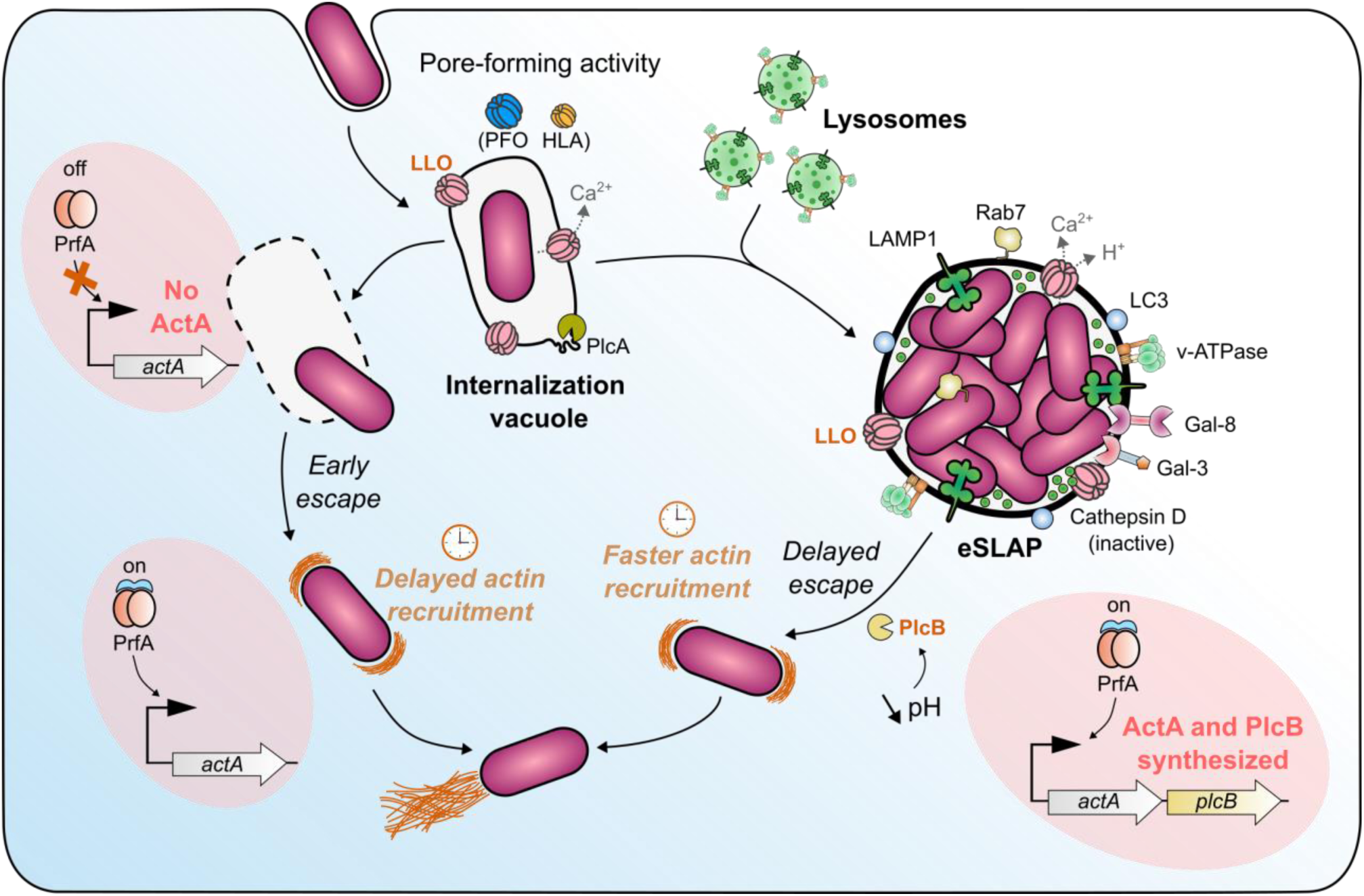
Extended model of eSLAP biogenesis and growth, and of *Listeria* virulence priming therein. In most cases, *Lm* rapidly escapes from the internalization vacuole into the cytosol. Alternatively, bacteria stay within an endomembrane compartment called eSLAP, where their replication and/or the enlargement of the vacuole require membrane permeabilization via the pore-forming activity of LLO. These compartments progressively grow and acquire late endosomal/lysosomal markers through interactions with host cell endomembrane trafficking pathways. Within eSLAPs, PrfA, the master regulator of *Lm* virulence genes, becomes activated, allowing the expression of the *actA-plcB* operon. The delayed rupture of eSLAP membranes is primarily mediated by the broad-range phospholipase PlcB, which requires not only PrfA-dependent expression but also pH-dependent maturation and release. Upon membrane breakdown, ActA-primed bacteria that escape from eSLAPs rapidly recruit actin, at a faster rate than those escaping early from internalization vacuoles.

### Endolysosomal maturation and growth of eSLAPs

While a variety of endomembrane compartments can in principle contribute to pathogen-containing vacuoles, our previous observation of the decoration of eSLAPs by Rab7 and LAMP1 pointed to an endolysosomal maturation route.^16^ The present study strengthens this conclusion by showing that eSLAPs are single-membraned and decorated by additional lysosolmal markers (LIMP2, ATP6V1A, and Cathepsin D), with no contribution of classical markers of the RE or Golgi apparatus. However, eSLAPs are not acidic and cathepsin D remains inactive, suggesting that these compartments are poorly degradative. In this respect, eSLAPs resemble the non-acidic, spacious single membrane-bound vacuoles that *Helicobacter pylori* and *Acinetobacter baumanii* can co-opt for survival or growth in epithelial cells.^43,44^ The analogy is reinforced by our TEM observations of structures reminiscent of the intraluminal vesicles found in late endosome MVBs, which were also observed in these two pathogen-containing vacuoles.

Mechanistically, we hypothesize that eSLAPs derive from the internalization vacuole by a succession of fusion events with lysosomes and/or late endosomes that accommodate bacterial growth by allowing the expansion of the vacuole membrane. Therein, *Lm* could also benefit from lysosomal degradation products as nutrients to sustain its replication. Importantly, the permeation of the vacuole membrane by a pore-forming toxin is a prerequisite to bacterial growth within eSLAPs, implying that when the membrane is intact, either the bacteria cannot access nutrients, or the fusion of internalization vacuoles with lysosomes is precluded. The fact that HLA can substitute for LLO in this permeation role indicates that the transit of small solutes such as ions is sufficient to support eSLAP enlargement. A plausible model is that the permeation of the vacuole membrane by LLO leads to local calcium efflux towards the cytosol, which in turn would drives the fusion of lysosomes with eSLAPs. This assumption stems from previous reports of calcium-dependent fusion events between lysosomes and phagosomes during the phagocytosis of *Salmonella* Typhimurium by macrophages or the growth of *Candida albicans* hyphae in phagosomes.^45–47^ Whether calcium is released from eSLAPs through LLO pores and modulates the fusion of lysosomes with the expanding vacuole will deserve special attention in future work aiming at identifying the molecular players involved in fusion events.

### Control of the balance between membrane permeation and disruption in eSLAPs

Our work has disentangled the roles in eSLAPs of two *Lm*-secreted effectors known to perturb the integrity of membranes: on the one hand, by permeating the vacuole membrane, the pore-forming activity of LLO is critical to support bacterial replication and/or vacuole expansion, while on the other hand, the phospholipase activity of PlcB is ultimately responsible for the breakdown of the vacuole membrane and the release of the bacteria in the cytosol. Our results do not exclude a possibility that once PlcB is activated, it acts in synergy with LLO in this release. Indeed, it was previously reported that the phospholipase activity of PlcB facilitated the cholesterol-dependent access of LLO to lipid bilayers and enhanced its capacity to disintegrate membranes.^48^ This differs from the mechanism of early escape from the internalization vacuole, which relies mostly on the activities of LLO and PlcA, as PlcB is not expressed at this stage.^18,22,25^

Consistently, our previous observation that the lifespan of eSLAPs was not correlated with LLO abundance suggested that the PFT alone had little impact on the stability of these compartments.^16^ Instead, our data support a model where LLO pore-forming activity results in a constant permeation of membranes, which we have discussed above as a requirement for eSLAP growth, and which is attested by the continuous recruitment of galectins. Permeated eSLAPs are non-acidic, which opposes the maturation of Mpl and of its substrate PlcB into their active forms. Although PrfA-dependent expression of the *actA-plcB* operon is induced by bacteria residing in eSLAPs, our data suggest that delayed maturation of PlcB favors the maintenance of these compartments. In this scenario, the lifespan of eSLAPs would be determined first by the timing of PrfA allosteric activation, and then by the probability that transient or moderate eSLAP acidification leads to the maturation of Mpl and PlcB in sufficient amounts to allow membrane rupture. The pores formed by LLO would likely slow-down PlcB maturation rate and contribute to eSLAP stability due to proton leakage towards the cytosol.

Interestingly, in macrophages, the pharmacological inhibition of PrfA allosteric transition has been shown to result in an abnormally high proliferation of *Lm* inside replication-permissive SLAP-like compartments (rSLAPs).^49^ Although the PrfA-dependent effector responsible for the escape from the precursors of these compartments in absence of treatment was not identified in this study, PlcB appears as a candidate of choice by analogy with our own data.

In the future, our model would benefit from a corroboration of the mechanistic evidence brought here by the use of genetics and pharmacological inhibitors with a live visualization of the molecular determinants of eSLAP rupture. For instance, the lifespan of eSLAPs could be correlated with PlcB abundance using a PlcB-FAST fusion, or with a precise monitoring of pH variations in eSLAPs using ratiometric probes.

### eSLAP, a niche where virulence is primed and a “bootcamp” preparing for cytosolic life

To date, most studies investigating the timing of allosteric activation of PrfA in human cells had concluded that it was triggered only once the bacteria gained cytosolic access. Elegant evidence for this model was brought by a screen for mutants allowing the intracellular survival of a “suicide” *Lm* strain that is killed when the promoter of *actA* is active.^28^ This screen yielded mutations targeting the capacity of *Lm* to exit the phagosome, emphasizing that phagosomal residence precluded PrfA activation. Nevertheless, it was more recently observed that a P*_actA_*-GFP construct could be induced by *Lm* within phagosomes in J774 macrophages, suggesting that PrfA activation can be primed in this compartment.^29^ In epithelial cells, to the best of our knowledge, the present work constitutes the first direct evidence of an induction of the *actA-plcB* operon in endomembrane compartments deriving from the internalization vacuole, and of the actual production of the ActA protein at this stage.

This induction results in the exposure of ActA at the surface of bacteria before they reach the cytosol, enabling the bacteria to recruit actin and initiate the polymerization of actin tails more rapidly upon escape from eSLAPs than when escaping internalization vacuoles. Additionally, it allows the production of PlcB which, after maturation, will lead to the disruption of eSLAPs. Being immediately motile upon reaching the cytosol can benefit the bacteria in several respects, which would deserve being assessed in the future. First, the bacteria could disseminate faster to neighboring cells, by analogy with the higher cell-to-cell spread rate observed when the interferon-dependent induction of IFITM3 inhibits the proteolysis of ActA in the phagosome.^29^ Faster cell-to-cell spread can amplify the penetration of bacteria within tissues or their shedding; it is also expected to promote bacterial fitness by bypassing nutrient limitation at the initial site of invasion.^50^ In addition, several studies have reported that the ActA-dependent recruitment of actin at the bacterial surface and cytosolic motility provided a protection of *Lm* against autophagy.^6,7,51^ Compared with an early exit from internalization vacuoles and immediate exposure to the cytosol, prolonged residence inside eSLAPs thus constitutes a transient niche where bacteria can replicate in an environment sheltered from cytosolic defenses, including inflammasomes, ubiquitin-ligases and xenophagy. In this “bootcamp”, the gradual priming of PrfA activity is likely to confer a second layer of protection from autophagy upon PlcB-dependent release in the cytosol, through ActA.

## Conclusion

Compared with the classical scenario of the intracellular life cycle of *Lm*, we are only starting to decipher the mechanisms that allow its prolonged vacuolar residence and replication, and to pinpoint some of its consequences with regards to the modulation of its virulence. The discovery that PrfA-dependent virulence is primed in eSLAPs and can prepare bacteria for a better success rate during the colonisation of the cytosol opens a window on the potential impact of avoiding an early exit from the internalisation vacuole. The fact that PlcB is gating the escape from eSLAPs also brings a rationale for the selection during evolution of the co-expression of *actA* and *plcB* in one operon, since it ensures that the bacteria able to escape eSLAPs are also equipped with ActA.

Yet, processes that may initially orient the fate of *Lm* towards a cytosolic or vacuolar life are still unknown. We are also far from reaching an integrated understanding of how eSLAPs or other intravacuolar niches may modulate the progression of infection *in vivo* or the efficiency of dissemination to new hosts. Although the coexistence of multiple routes is likely to alter the dynamics of bacterial progression in tissues, the sensing of bacteria by immune surveillance mechanisms or their sensitivity to immune responses, at this stage we lack robust models that would recapitulate the three-dimensional structure of tissues while allowing the tracking of individual bacteria and of their outcomes following relatively rare events such as eSLAPs over prolonged time lapses. Identifying the tissues and organs where eSLAPs are the most relevant would represent a first step in this direction.

## Materials and Methods

### Bacterial strains, plasmids and culture conditions

The bacterial source strains used for this work (Table S1) were *Listeria monocytogenes* LL195 as a main infection model. For plasmid cloning, *Escherichia coli* NEB5α (New England BioLabs) and *Escherichia coli* NEBStable (New England BioLabs) for lentiviral vector cloning were used. All strains were grown at 37 °C under shaking at 180 rpm in Luria Bertani (LB) medium for *E. coli* (except *E. coli* NEBStable that was grown at 30 °C), in brain heart infusion (BHI) or *Listeria* synthetic medium (LSM)^52^ for *Lm*. Whenever required, media were supplemented with antibiotics for plasmid selection (kanamycin, 30 μg/μL, ampicillin, 100 μg/μL, chloramphenicol, 20 μg/μL for *E. coli* and 7 μg/μL for *Lm*).

For allelic replacements in *Lm*, 500 to 1,000 bp homology regions flanking the genomic sequence to be deleted or inserted were PCR-amplified using the primers listed in Table S2. *Lm* genes were amplified from the LL195 genome. The sequences of *pfo* and *hla* were respectively amplified from the plasmids pTrcHisA-*pfo*^53^ and from pFHA-*hla* (gift from Lionel Navarro). The resulting PCR products were assembled into the temperature-sensitive shuttle vector pMAD by Gibson assembly between the BglII and SalI restriction sites. Allelic exchange was carried out as previously described.^54^ For constitutive mCherry expression from an inert locus, a P_HYPER_-mCherry cassette was integrated in the intergenic region between genes CCO63057 and CCO63058 in the LL195 genome (corresponding to *lmo0439* and *lmo0440* in EGD-e) via pMAD-driven allelic exchange. No difference in bacterial entry or intracellular replication rates was identified in this strain compared to the non-modified LL195 strain.

The PCR-amplified constructs with the P*_actA_* promoter were integrated at the SmaI-SalI sites of pAD_2_-P*_inlC_*-GFP.^55^ All other constructs under control of the constitutive P_HYPER_ promoter were integrated at the EagI-SalI sites of pAD_1_-cGFP.^55^ All pAD-derived plasmids were integrated in the genome of *Lm* LL195 at the *tRNA^Arg^* locus by electroporation^56^, except pHpPL3-mCherry and pAD_1_-cGFP plasmids that were introduced by conjugation.

The PCR amplified products for lentivirus production were integrated at the EcoRI-BamHI sites of pLV-EF1α-IRES-Puro.^57^ In order to favor the expression of EYFP-CBD_500_ in LoVo cells, the DNA coding sequence for CBD_500_ was codon-optimized for *Homo sapiens*. The optimized sequence was obtained as a synthetic Gene Fragment (Eurofins genomics) (Table S2).

### Culture of cell lines

LoVo cells, a human intestinal epithelial cell line originating from a colon adenocarcinoma (male, ATCC CCL-229) were used as a main host cell model. Cells were routinely maintained in their respective formulated media, supplemented with 10 % heat-inactivated fetal bovine serum (cat#p30-3306, PANBiotech) without antibiotics in a humidified incubator at 37 °C and 5 % CO_2_. Formulated media were: for LoVo, DMEM, low glucose, GlutaMAX supplement, pyruvate medium (cat#21885025, ThermoFisher Scientific); for Hep G2, Huh-7, LS174T and MCF-7, Eagles’ MEM (cat#10009CV, Corning); for HCT-116 and SK-BR-3, McCoy’s 5A medium (cat#16600082, ThermoFisher Scientific), for T84, DMEM/F-12 (cat#11320033, ThermoFisher Scientific).

### Lentivirus production and generation of polyclonal stable cell lines

All cell lines expressing fusions of a fluorescent marker with human proteins or with CBD_500_ were obtained by lentiviral transduction. 1,2 x 10^6^ HEK293 FT cells were plated in a 60 mm dish. 24h later, cells were transfected with 5.5 μg of total DNA comprising the vector plasmid with the desired transgene (pLV-EF1a-IRES-Puro), the packaging plasmid psPAX2 and the envelope plasmid pCMV-VSVG (Table S1) at a molar ratio of 4:3:1 using Lipofectamine 3000 according to the manufacturer’s protocol (L3000008, Thermofisher Scientific). 36 h post-transfection, 4 mL of the supernatant containing the lentiviruses were filtered through a 0.45-μm filter pore. 1 × 10^6^ LoVo cells and polybrene (TR-1003, Sigma-Aldrich; 8 μg/mL) were added to the supernatant and plated in 10-cm dishes. The infection was carried out for 6 h after which the medium was renewed. 24 h post infection, transduced cells were selected by adding puromycin (1 μg/mL) until 100 % cell death was observed in the control (non-transduced) cells.

### Precipitation of bacterial secreted proteins

Protein from *Lm* strains were purified as previously described.^58^ Overnight cultures of *Lm* grown in LSM or LSM + 10 mM glutathione (GSH) (for PrfA induction) were centrifuged at 6,000 × *g* for 1 min at 4 °C. The recovered supernatants were filtered through a 0.22-μm pore filter. Sodium deoxycholate (0.2 mg/mL) and PMSF (0.5 mM) were added to the filtered fractions and incubated on ice for 30 min. Trichloroacetic acid (TCA) was added at a final concentration of 12 %, then the mix was incubated at 4 °C on a rotating wheel for 1 h 30. The precipitated proteins were pelleted by centrifugation for 15 min at 16,000 × *g* at 4 °C, then washed twice in ice-cold acetone. The protein pellets were air-dried 5 min at 20 °C, dissolved in SB 1,5 X and denatured for 5 min at 95 °C prior to protein analysis.

### Hemolysis assay of *Listeria* strains

The supernatant of ON cultures of the different *Lm* strains grown in BHI were recovered by centrifugation for 1 min at 6,000 × *g* followed by filtration through a 0.22-μm filter. Serial two-fold dilutions of the supernatants using PBS pH 5.6 with 0.1 % BSA were done in a 96-well plate. Erythrocytes from defibrinated horse blood (CL1500-50D, Cedarlane Laboratories Limited) were washed twice in PBS pH 6.4 and diluted 1:10 in PBS pH 5.6, then 50 μL of the dilution was added in each well. Plates were incubated at 37 °C for 30 min before centrifugation for 10 min at 400 × *g*, then the hemolytic titers were calculated as the reciprocal of the dilution for which 50 % of the hemolysis was observed.^59^ Hemolysis titers were normalized with the supernatant of the WT strain.

### Protein analysis

Equal amounts of protein were separated on a gel by SDS-PAGE. For direct staining of total proteins, the gel was stained with Coomassie Brilliant blue G-250 as described before.^16^ For western blot analysis, proteins were transferred on a nitrocellulose membrane (Amersham) using BioRad wet transfer tank for 1 h at 100 V. The membrane was saturated in blocking buffer (5 % skimmed milk and 0.05 % Tween-20 in TBS) for 30 min at 20 °C. The membrane was then incubated with the appropriate primary antibody in blocking buffer ON at 4 °C followed by incubation with anti-mouse or rabbit IgGs coupled to HRP, (cat#A120101P and A90-116P respectively, Bethyl) at 1:50,000 in the same buffer. Proteins were revealed using SuperSignal West Pico PLUS (cat #34578, Thermo Scientific) and imaged with a Las4000 imager (Amersham). The primary antibodies and respective dilutions used were the following: rabbit anti-LLO (R176)^60^ 1:10,000; anti-InlC (R117)^61^ 1:1,000; anti-EF-Tu (R114)^62^ 1:20,000; anti-Gal8 (cat#HPA030491, Atlas Antibodies) 1:1,000; anti-Gal3 (clone D4I2R cat#87985 Cell Signaling Technology) 1:1,000; anti-PABPC1 (cat#HPA045423 Atlas Antibodies) 1:1,000; and mouse anti-βtubulin (cat#BLE903401 BioLegend) 1:2,000.

### Cell infection, immunofluorescence and microscopy of infected cells

85,000 LoVo cells were seeded 48 h before infection in 24-well plates with glass coverslips. Cells were infected with late exponential/early stationary *Lm* culture (OD_600nm_ between 1,5 and 2) at the desired multiplicity of infection (MOI) and the bacteria were synchronously brought in contact with the cell monolayer by centrifugation at 200 × *g* for 1 min. When indicated, bafilomycin A1 (100 nM) or monensin (100 μM) were added to infected cells and incubated only during the first hour of infection. At 1 hpi, cells were washed twice with DMEM with 50 μg/mL gentamicin and medium was replaced by DMEM with FBS 10 % and gentamicin 25 μg/mL until the end of the infection. Cells were then washed in PBS and fixed in PFA 4 % in PBS for 20 min at 20 °C or fixed/permeabilized with ice-cold 100 % methanol at 4 °C for 10 min, for detection of ATP6V1A, Cathepsin D and LIMP-2. Cells were washed twice and permeabilized with Triton 0,5 % for 5 min at 20 °C. Cells were blocked in blocking buffer (5 % BSA and 0,05% Tween-20 in PBS) at 4 °C overnight. Then, coverslips were incubated with the primary antibody in blocking buffer for 1 h at 20 °C in the dark. Coverslips were washed three times in large amounts of PBS then incubated with the appropriate secondary antibody (Alexa Fluor 647 or 488-conjugated secondary anti-rabbit antibodies, respectively cat# A21245 and cat# A11008, Molecular probes, at a 1:500 dilution), and when relevant, other fluorescent dyes (Acti-stain 647 phalloidin #PHDN1, Cytoskeleton, 70 nM and DAPI, 0.1 μg/μl or Hoechst 33342, 1 μg/μl) for 45 min in the dark. The primary antibodies and dilutions used were the following: anti-rabbit against ActA (P473)^63^ 1:500; anti-rabbit from GeneTex against ATP6V1A (GTX110815) 1:400; anti-rabbit from Cell Signalling Technologies against cathepsin D (clone E179 cat #69854), 1:200 LIMP-2/SCARB2 (clone E2Z5F cat#27960) 1:200 and LAMP1 (clone D2D11 cat#9091) 1:200. Coverslips were washed twice in PBS and once in sterile water, then mounted on a glass slide with Fluoromount-G. Infected cells were observed with a Nikon Ti epifluorescence microscope (Nikon), connected to a digital CMOS camera (Orca Flash 4.0, Hamamatsu). Illumination was achieved using a SOLA-SE 365 source (Lumencor) and the following excitation/emission/dichroic filter sets (Semrock): DAPI or Hoechst, 377(50)/447(60)/FF409-Di03; GFP or Alexa 488, 472(30)/520(35)/FF495-Di03; mCherry, 562(40)/632(22)/dic FF562-Di03; Alexa 647 or Cy5, 630(30)/684(24)/dic FF655-Di01. Images were acquired with Nikon apochromat 60x objective lens (NA 1.4). Image acquisition and microscope control were actuated with the μManager software (RRID:SCR_016865), and processed with Fiji. Each picture is representative of the infected cell population.

### Gentamicin protection assay

A classical gentamicin protection assay was used to evaluate bacterial entry in cells. Cells were infected with *Lm* strains as described above, except the gentamicin wash step was carried out at 30 min post-infection. The infection was continued for another 30 min, then cells were washed once in PBS and lysed in ice-cold sterile water by pipetting up and down. Serial dilutions were plated on LB plates and incubated at 37 °C for 24 h. The colony-forming units (cfu) were counted and normalized to cell counts to determine the number of bacteria per cell.

### Live microscopy of infected cells

At the appropriate time post infection, infected cells were observed in DMEM without phenol red supplemented with 25 μg/mL gentamicin, 500 nM of the appropriate SiR-dye: SiR-Lysosome (SC012, Spirochrome) or SPY650-Fast-Act (SC505, Spirochrome), and when relevant, the fluorogenic chromophore 5 μM of HBR-3,5DM ((*Z*)-5-(4-Hydroxy-3,5-dimethylbenzylidene)-2-thioxothiazolidin-4-one) for visualization of FAST2-producing *Lm* strains. Samples were imaged with a Nikon Ti PFS microscope coupled to a spinning disk confocal device (CSU-XI-A1, Yokogawa), connected to the camera ORCA Flash 4.0 LT (Hammatsu) and equipped with a cube for temperature control and a brick gas mixed for CO_2_ and humidity control (Life Imaging Services). Image acquisition was performed with the MetaMorph software (Molecular Devices, RRID:SCR_002368). Fluorescence was triggered by three lasers at 491 nm wavelength for GFP, eYFP and mAG, 561 nm for mCherry and mRFP, and 635 nm for SiR-Lysosome or SPY650-Fast-Act. Images were acquired with apochromat 60x objective lens (NA 1.4) in 0.7 μm step-*z*-stacks every 5 to 10 min. Acquisition parameters were similar for all samples of an experiment. Maximum intensity signals of the *z*-stack were integrated with Fiji.

### Quantification of eSLAP frequency in infected cells

For eSLAP quantification, cells were infected with the appropriate *Lm* strains expressing mCherry for 3 h, then processed for immunofluorescence staining of LAMP1 with Alexa 488, labelling of F-actin with Acti-stain 647 phalloidin, and DAPI-staining of the cell nuclei as described above. For each condition and each replicate, 16 to 20 random fields were selected using the actin channel alone to avoid unconscious user bias during image acquisition, then images were acquired in all fluorescence channels. Images were processed using a custom Python script (available upon request), which quantified the number of nuclei per field using the DAPI channel —used as a proxy for the number of cells—, the number of bacteria per cell based on mCherry fluorescence, and identified eSLAPs by detecting spherical clusters of at least three bacteria colocalizing with LAMP1 signal. In more detail, Otsu thresholding was first performed on the mCherry, Alexa 488 and DAPI channels to segment respectively the bacteria, LAMP1-stained compartments, and cell nuclei. From the mask corresponding to bacteria, an intermediate mask was created that contained only bacterial aggregates, by excluding objects for which the surface was below three times the average surface of one bacterium. The eSLAP mask was then obtained as the intersection of the LAMP1 mask and of the intermediate mask for bacterial aggregates. The numbers of eSLAPs, of bacteria and of nuclei per image were quantified using the *label* function of the package *skimage* (<u>scikit-image 0.20.0</u>). For eSLAPs, detected objects were filtered with criteria on (*i*) area (surface above three bacteria), (*ii*) solidity, to evaluate shape regularity and select objects with smooth edges, and (*iii*) eccentricity, to evaluate the overall shape of objects and select the ones that were rounded or slightly oval-shaped. The total number of bacteria was evaluated by dividing the total area of the mask for bacteria by the average surface of one bacterium. Nuclei were only filtered with a criterium on their area. The output for each image was exported in a .csv file containing the image name, the counts for nuclei, eSLAPs and bacteria, the number of bacteria per cell, and the number of eSLAPs per 100 cells.

Before running automated quantifications, the script was validated by comparing its outputs with results of manual quantifications performed by an independent user on 30 images for nuclei (three biological replicates, 681 nuclei) and 36 images for eSLAPs (two biological replicates, 68 eSLAPs). 92% of nuclei were accurately detected by the script, and 8% of nuclei were not detected (typically nuclei that were too close to each other and were classified as a single nucleus). For eSLAPs, 95 % of the detected objects detected by the script were also identified as eSLAPs by manual annotation, while 5 % appeared as false positives. 6% of the eSLAPs counted by the user in the images were not detected by the script.

The eSLAP frequency for each image was calculated by normalizing the number of eSLAP/100 cells by the mean value of eSLAP/100 cells of the corresponding control condition. Data are represented in “SuperPlots” as described in Lord *et al*. where each biological replicate is symbol and color-coded.^64^

### Quantification of Galectin fluorescence

LoVo cells expressing mAG-Gal3 or EGFP-Gal8 were infected and imaged as indicated above. Images from movies were *z*-projected by maximum intensity. Fluorescence of mAG (Gal3) and EGFP (Gal8) was measured as the mean value of pixel intensity in a Region Of Interest (ROI) manually positioned over the eSLAP for each time point until the dispersion of bacteria was observed, corresponding to eSLAP rupture. The background cytosolic fluorescence intensity of Gal3 and Gal8 was also measured at each time point in a ROI positioned in the cytosol of the same cell, in a region devoid of bacteria. The background mean cytosolic fluorescence intensity was subtracted from the eSLAP-associated intensity at each corresponding time point.

### Quantification of PrfA activation in eSLAPs

Cells were infected with *Lm* mCherry [pAD-P*_actA_*-*gfpmut2*] for 25 min or 3 h and processed for immunofluorescence using an anti-ActA antibody as described above. PrfA activation was identified based on GFP fluorescence, and the presence of ActA was assessed by immunolabeling. ActA-positive and GFP-positive eSLAPs were manually quantified, with approximately 100 eSLAPs analyzed per replicate.

### Measurement of *Lm* actin recruitment timing

LoVo cells expressing EYFP-CBD_500_ were infected as described above with *Lm* expressing mCherry at MOI 20, except SPY650-FastAct was additionally added at a final concentration of 500 nM, 2 h before imaging. For the assessment of actin recruitment by bacteria escaping the internalization vacuole, the imaging was started at 30 min post-infection. Bacteria escaping eSLAPs were started being imaged at 4 hpi. Images were acquired in 0.5 μm step-*z*-stacks every 2.5 min, and acquisition parameters were similar for all samples of an experiment. Maximum intensity signals of the *z*-stack were integrated with Fiji. Actin recruitment time was manually determined and corresponded to the interval between the acquisition of the EYFP-CBD_500_ signal around the bacterium, indicating bacterial contact with the cytosol, and the appearance of the actin coat surrounding the bacterium. Only bacteria for which vacuole rupture was clearly visible were included in the analysis. To be considered, bacteria not labelled by EYFP-CBD_500_ had to be already present in the field of view at least one time point before the detection of the EYFP-CBD_500_ signal, to ensure their presence within a vacuole at that moment. It was observed that some bacteria residing within eSLAPs exhibited a faint EYFP-CBD_500_ labeling prior to rupture, likely reflecting the permeabilized nature of eSLAP compartments as observed with galectin staining. In such cases, the rupture time was defined as the time point at which the mean EYFP-CBD_500_ signal intensity within the eSLAP reached at least half the mean signal measured for a cytosolic bacterium.

### Transmission electron microscopy of infected cells

Infected cells were washed once in PBS, then fixed with 2.5 % glutaraldehyde in PBS for 1 h at 20 °C. Cells were washed three times in PBS then post-fixed 30 min in 1 % osmium tetroxide on ice with agitation. Cells were washed twice in PBS, then gradually dehydrated in ethanol solutions of increasing concentrations (25 %, 50 %, 70 %, 5 min each) before contrast staining in methanol 70 % containing 1 % uranyl acetate for 30 min at 20 °C. Dehydration was then continued in ethanol solutions (90 to 100 %). Samples were infiltrated with graded concentrations of araldite 502 resin in ethanol (30 %, 50 %, 70 % and 100 %, 5 min each), then embedded in araldite 502. Polymerization was carried out for 48 h at 60 °C. Ultrathin sections (70 nm) were obtained with an ultramicrotome (Ultracut UC6, Leica). Observations were performed with a Transmission Electron Microscope TECNAI 12BT at 80 kV (Thermofisher/FEI) equipped with a CCD Orius 1000 camera (Gatan). Image acquisition was performed with DigitalMicrograph.

### Statistical analysis

All analyses were performed with GraphPad Prism 9. All data sets were tested for normality with Shapiro-Wilkinson test and homoscedasticity was assumed. For normally distributed data sets, differences in the means between two groups was assessed by an unpaired two-tailed Student’s *t*-test. When data were not normally distributed, a Mann-Whitney *U* test was applied. For comparison involving more than two groups, a one-way ANOVA followed by Dunnett’s multiple comparisons test was used, while a Kruskal-Wallis followed by Dunn’s multiple comparisons test was performed for data that did not fit a normal distribution. When two independent variables were involved, a two-way ANOVA with Šídák’s correction for multiple comparison was used. A *p*-value less than 0.05 was considered statistically significant.

## Acknowledgements

We are grateful to Vinko Besic for generating the Δ*plcA* and Δ*plcB* deletion mutants in the LL195 background, to Caroline Peron-Cane for advice with live microscopy, to Philéas Larcher for feedback on the manuscript, and to the IBENS imaging facility IMACHEM.

For the purpose of Open Access, a CC-BY public copyright licence has been applied by the authors to the present document and will be applied to all subsequent versions up to the Author Accepted Manuscript arising from this submission.

## Author contributions

AL and TP designed the project, devised experiments and interpreted results. TP performed most experiments with the participation of SGM, AV and AC. TP, SGM and ND analyzed image data. TP and AL wrote the manuscript.

## Supplemental information

Supplemental information to this work includes supplemental Fig S1 to S3, supplemental Movies S1 to S14, and supplemental Tables S1 and S2.

**Fig S1.**
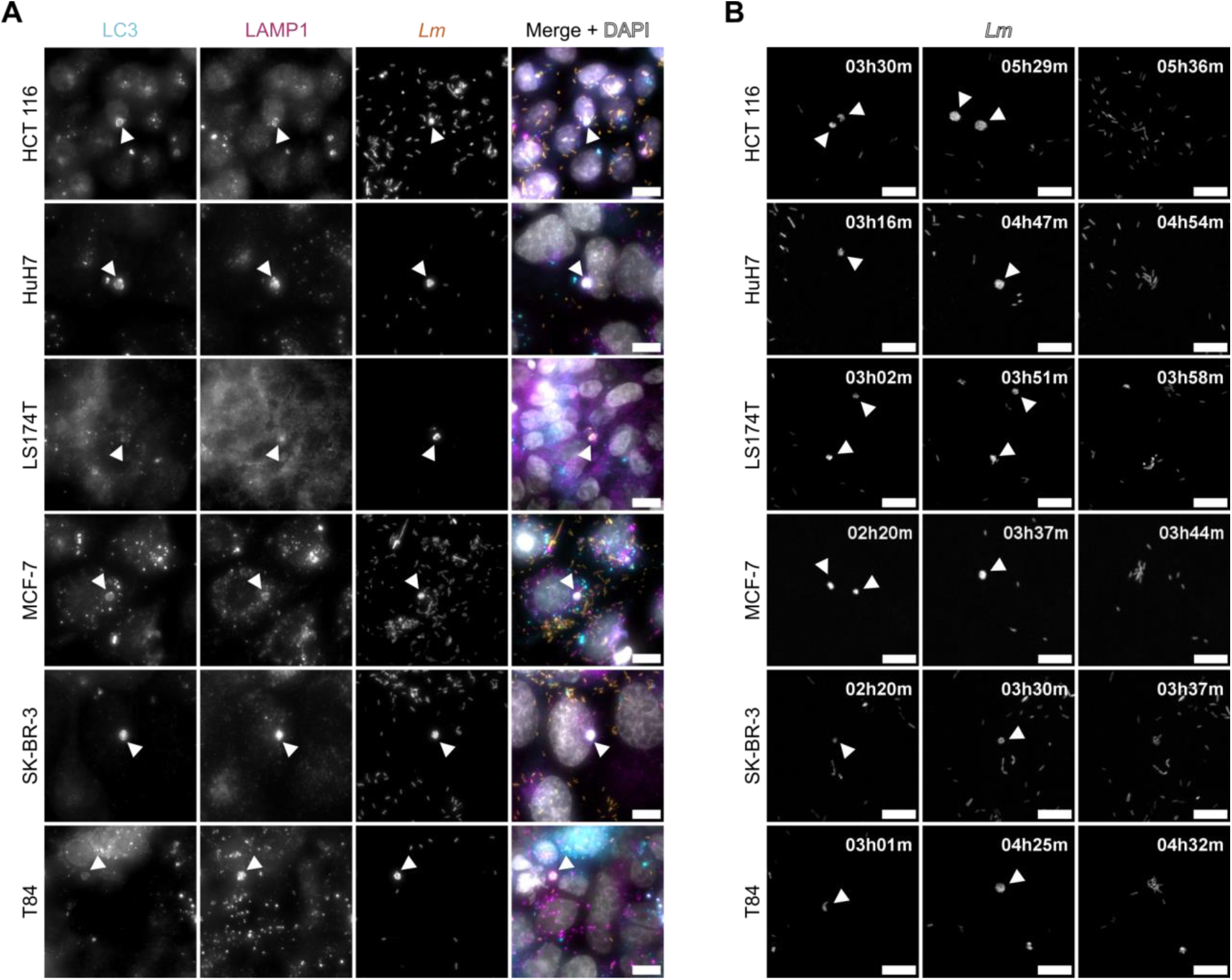
eSLAP occurrence in other cancer epithelial cell lines. Different cell lines were infected with *Lm* WT expressing mCherry at (A) MOI 50 (HCT 116, Huh-7, LS174T, SK-BR-3, T84) or MOI 100 (MCF-7) and (B) MOI 5 (SK-BR-3), MOI 10 (HCT 116, MCF-7) or MOI 20 (Huh-7, LS174T, T84). (A) Immunofluorescence staining against LC3 (cyan) and LAMP1 (magenta) at 3 hpi. *Lm* mCherry is shown in orange. Nuclei stained by DAPI are displayed in grey. (B) Representative live spinning disk microscopy images of the infected cell lines at different time points showing the replication of *Lm* mCherry (in grey) in eSLAPs. White arrowheads point (A) at eSLAPs and the associated markers and (B) at the growing eSLAPs. Scale bars: 10 μm.

**Fig S2.**
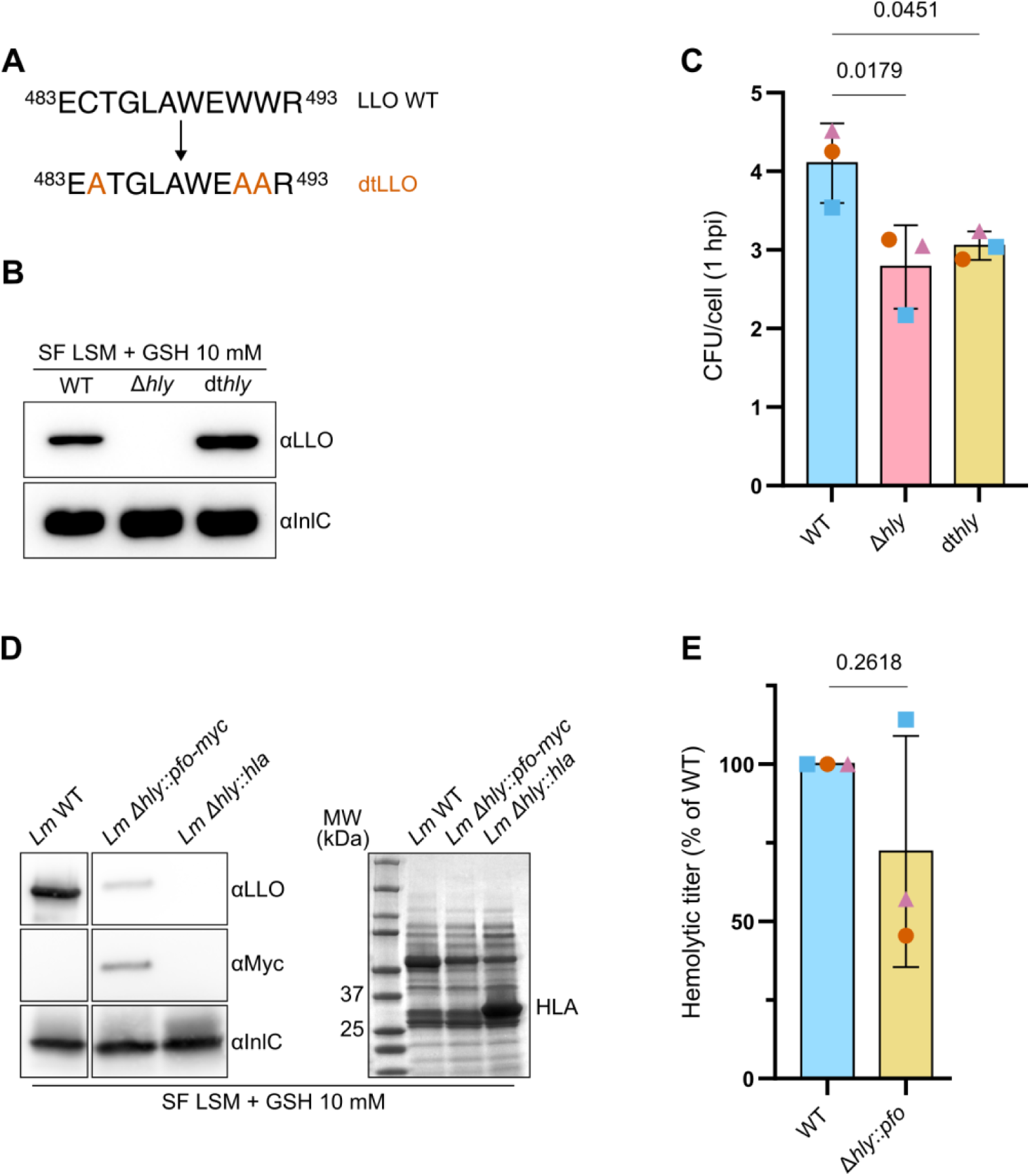
Secretion and hemolytic properties of pore-forming toxins. (A) Diagram showing the characteristic hendecapeptide sequence of LLO (amino acid 483 to 493) and the three mutated amino acids (in orange) in the detoxified LLO (dtLLO). (B) The secreted fractions (SF) from overnight cultures of *Lm* WT, *Lm* Δ*hly*; and *Lm* dt*hly* in LSM + GSH 10 mM were analyzed by immunoblot with antibodies against LLO, with InlC used as a loading control. (C) Assessment of the differential invasiveness of the three strains by counting the number of CFU per cell after gentamicin protection assays in LoVo cells infected for 1 h at a MOI of 100. (D) Secreted proteins from overnight cultures of *Lm* WT, *Lm* Δ*hly::pfo*-*myc*, and *Lm* Δ*hly::hla* in LSM + GSH 10 mM were analyzed by immunoblot with antibodies against LLO and Myc, with InlC used as a loading control (left) or by Coomassie staining (right). Due to cross-reactivity, the antibody against LLO faintly detects PFO-Myc, with a shifted size compared to that of LLO. On the Coomassie stain, the band corresponding to α-hemolysin (HLA) appears between 37 and 25 kDa. MW: molecular weight. (E) The hemolytic titer of *Lm* secreting LLO (WT) or PFO (Δ*hly*::*pfo*) was quantified on horse erythrocytes, and normalized to the WT strain taken as a reference. For (C, E), the plot represents the mean ± SD of three independent experiments, with data from independent experiments being displayed with three different symbols and colors. The *p*-values indicate the results of a one-way ANOVA with Dunnett’s multiple comparison test in (C), and of a two-tailed Student’s *t*-test in (E) assuming equal variances.

**Fig S3.**
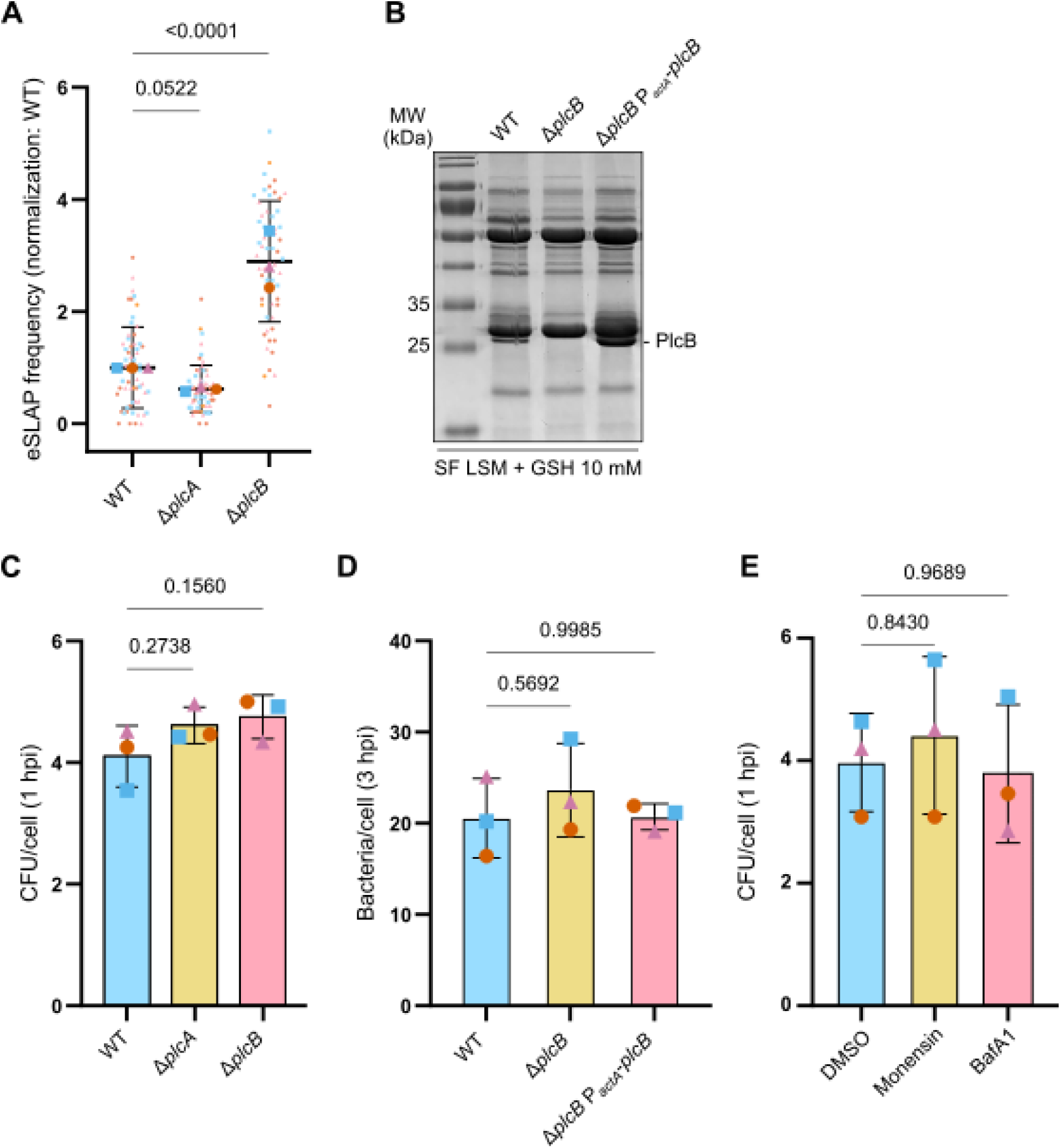
Effects of phospholipases, bafilomycin and monensin on the entry, replication and eSLAP frequency of *Lm* in LoVo cells. (A) Quantification of eSLAP frequency in LoVo cells infected for 3 h by *Lm* WT, Δ*plcA* and Δ*plcB* strains (MOI 100), all expressing mCherry. (A) Colloidal Coomassie staining of the secreted proteins from overnight cultures in *Listeria* synthetic medium (LSM) + glutathione (GSH, 10 mM) of *Lm* WT, Δ*plcB* and Δ*plcB*-P*_actA_*-*plcB*. The position of the band corresponding to PlcB is indicated, highlighting its expression by the WT and complemented strains. (C) Assessment of the differential invasiveness of the strains *Lm* WT, Δ*plcA* and Δ*plcB* by counting the number of CFU per cell after gentamicin protection assays in LoVo cells infected for 1 h at MOI 100. (D) Quantification of the number of bacteria per cell from microscopy slides of the experiment shown in Fig 4A, corresponding to 3 h of infection of LoVo cells with Lm WT, Δ*plcB* or complemented for *plcB* expression, all expressing mCherry. (E) Assessment of *Lm* entry efficiency by gentamicin protection assay on LoVo cells treated with DMSO vehicle, monensin (100 μM) or bafilomycin A1 (BafA1, 100 nM), and infected by *Lm* WT at MOI 100 for 1 h. (A, C, D and E) Data from three independent experiments are displayed with three different symbols and colors. The plots represent the mean ± SD. The *p*-values indicate the results of one-way ANOVAs with Dunnett’s correction for multiple comparisons, assuming equal variance.

**Movie S1. Observation of an eSLAP in HCT 116 cells infected by *Lm*.**

HCT 116 cells infected with *Lm* expressing mCherry (orange) were observed between 3 h 30 and 7 h post-infection by spinning disk microscopy. Actin is shown in magenta. Scale bar, 10 μm.

**Movie S2. Observation of an eSLAP in Huh-7 cells infected by *Lm*.**

Huh-7 cells infected with *Lm* expressing mCherry (orange) were observed between 3 h 15 and 7 h 45 post-infection by spinning disk microscopy. Actin is shown in magenta. Scale bar, 10 μm.

**Movie S3. Observation of an eSLAP in LS174T cells infected by *Lm*.**

LS174T cells infected with *Lm* expressing mCherry (orange) were observed between 3 h and 6 h 30 post-infection by spinning disk microscopy. Actin is shown in magenta. Scale bar, 10 μm.

**Movie S4. Observation of an eSLAP in MCF-7 cells infected by *Lm*.**

MCF-7 cells infected with *Lm* expressing mCherry (orange) were observed between 2 h 20 and 6 h 45 post-infection by spinning disk microscopy. Actin is shown in magenta. Scale bar, 10 μm.

**Movie S5. Observation of an eSLAP in SK-BR-3 cells infected by *Lm*.**

SK-BR-3 cells infected with *Lm* expressing mCherry (orange) were observed between 2 h 20 and 7 h 15 post-infection by spinning disk microscopy. Actin is shown in magenta. Scale bar, 10 μm.

**Movie S6. Observation of an eSLAP in T84 cells infected by *Lm*.**

T84 cells infected with *Lm* expressing mCherry (orange) were observed between 3 h and 6 h 30 post-infection by spinning disk microscopy. Actin is shown in magenta. Scale bar, 10 μm.

**Movie S7. Observation of the decoration of eSLAPs by mAG-Gal3 in cells infected by *Lm*.**

LoVo cells stably expressing mAF-Gal3 (cyan) infected with *Lm* expressing mCherry (orange) were observed between 3 h 45 and 7 h 45 post-infection by spinning disk microscopy. Scale bar, 10 μm.

**Movie S8. Observation of the decoration of eSLAPs by EGFP-Gal8 in cells infected by *Lm*.**

LoVo cells stably expressing EGFP-Gal8 (cyan) infected with *Lm* expressing mCherry (orange) were observed between 2 h 45 and 7 h 45 post-infection by spinning disk microscopy. Scale bar, 10 μm.

**Movie S9. Imaging of the replication of *Lm* WT GFP within eSLAPs in LoVo cells expressing mRFP-LC3.**

LoVo cells stably expressing mRFP-LC3 (cyan) infected with *Lm* WT expressing GFP (orange) were observed between 3 h and 7 h 30 post-infection by spinning disk microscopy. Actin is shown in magenta. Scale bar, 10 μm.

**Movie S10. Imaging of the replication of *Lm* Δ*hly::pfo-myc* GFP within eSLAPs in LoVo cells expressing mRFP-LC3.**

LoVo cells stably expressing mRFP-LC3 (cyan) infected with *Lm* Δ*hly::pfo-myc* expressing GFP (orange) were observed between 3 h and 7 h 30 post-infection by spinning disk microscopy. Actin is shown in magenta. Scale bar, 10 μm.

**Movie S11. Imaging of the replication of *Lm* Δ*hly::hla* GFP within eSLAPs in LoVo cells expressing mRFP-LC3.**

LoVo cells stably expressing mRFP-LC3 (cyan) infected with *Lm* Δ*hly::hla* expressing GFP (orange) were observed between 3h and 7h30 post-infection by spinning disk microscopy. Actin is shown in magenta. Scale bar, 10 μm.

**Movie S12. Imaging of the replication of *Lm mCherry* P*_actA_*-*actA-fast2* within eSLAPs in LoVo cells.**

LoVo cells infected with *Lm* expressing constitutively mCherry (magenta) and *actA-fast2* from its natural promoter (orange) were observed between 4 h and 6 h 15 post-infection by spinning disk microscopy. Scale bar, 10 μm.

**Movie S13. Imaging of the labelling by EYFP-CBD_500_ and SPY-FastAct of bacteria escaping the internalization vacuole.**

LoVo cells stably expressing EYFP-CBD_500_ (magenta) infected with *Lm* expressing mCherry (orange) were observed between 30 min and 2h35 post-infection by spinning disk microscopy. Actin is shown in cyan. Scale bar, 10 μm.

**Movie S14. Imaging of the labelling by EYFP-CBD_500_ and SPY-FastAct of bacteria escaping eSLAPs.**

LoVo cells stably expressing EYFP-CBD_500_ (magenta) infected with *Lm* expressing mCherry (orange) were observed between 4h30 and 7h05 post-infection by spinning disk microscopy. Actin is shown in cyan. Scale bar, 10 μm.

**Table S1. Bacterial strains and plasmids.**

**Table S2. Oligonucleotides and gene fragments.**

